# An RNA degradation complex required for silencing of Polycomb target genes

**DOI:** 10.1101/2019.12.23.887547

**Authors:** Haining Zhou, Gergana Shipkovenska, Marian Kalocsay, Jiuchun Zhang, Zhenhua Luo, Steven P. Gygi, Danesh Moazed

## Abstract

Polycomb Repressive Complex (PRC) 1 and 2 are histone-modifying and chromatin-binding complexes that are required for silencing of developmental regulatory genes and genes that control cellular proliferation. Their gene silencing functions are thought to involve chromatin compaction and condensate formation but whether other mechanisms contribute to silencing is unknown. Here we show that the rixosome, a conserved RNA degradation complex with roles in ribosomal RNA processing and heterochromatic RNA degradation in fission yeast, associates with human PRC complexes, is recruited to promoters of Polycomb target genes in differentiated cell lines and embryonic stem cells, and is required for efficient silencing of Polycomb target genes. These findings reveal an unanticipated role for RNA degradation in Polycomb-mediated silencing.

## Introduction

The Polycomb group genes play important roles in silencing of cell type-specific genes outside of their proper domains of expression in flies and mammals (Schuettengruber et al., 2017, Simon and Kingston, 2013, Margueron and Reinberg, 2011). The mechanism of Polycomb-mediated silencing involves the tri-methylation of histone H3 lysine 27 (H3K27me3) by the EZH1 or EZH2 subunit of PRC2 and mono-ubiquitylation of histone H2A lysine 119 (H2AK119ub1) by the RING finger E3 ubiquitin ligase subunit of PRC1. These modifications orchestrate a series of regulatory events, including allosteric activation of the methyltransferase activity of EZH2 by binding of EED to H3K27me3 and crosstalk between the PRC1 and PRC2 complexes through subunits in each complex that recognize the modification that the complex itself catalyzes as well as the one catalyzed by the other complex (Simon and Kingston, 2013, Margueron and Reinberg, 2011). These recognition events are not only required for the epigenetic inheritance of Polycomb silencing, but also play critical roles in the silencing functions of these complexes. For example, H3K27me3 provides a binding site for the PRC1 chromodomain CBX2 subunit, which promotes the compaction of nucleosome arrays and has more recently been found to mediate liquid-liquid phase separation (LLPS)(Schuettengruber et al., 2017, Francis et al., 2004, Margueron et al., 2008, Terranova et al., 2008, Lau et al., 2017, Plys et al., 2019, Tatavosian et al., 2019). It has therefore generally been thought that the mechanism of silencing involves the exclusion of RNA polymerase II via compaction or condensate formation. However, previous studies also provide evidence for the presence of the general transcription machinery and RNA Polymerase II (Pol II) at promoters of Polycomb repressed genes (Schuettengruber et al., 2017, Breiling et al., 2001, Dellino et al., 2004). The mechanisms that prevent Pol II from productive transcription of Polycomb targets remains unknown.

The rixosome is a highly conserved and essential multienzyme complex with RNA endonuclease and polynucleotide kinase activities required for rRNA processing and ribosome biogenesis (Fromm et al., 2017). We recently showed that, in the fission yeast *Schizosaccharomyces pombe*, the rixosome also plays a critical role in the spreading of histone H3 lysine 9 (H3K9) methylation into actively transcribed regions and is required for epigenetic inheritance of heterochromatin (Shipkovenska et al., 2019). To test whether the human rixosome plays similar roles in heterochromatin regulation, we purified the complex from human cells and analyzed its composition by mass spectrometry. We show that in human embryonic stem cells and differentiated cell lines the rixosome associates with both PRC1 and PRC2 complexes, and is recruited to Polycomb target genes where it promotes degradation of nascent transcripts and release of RNA polymerase II. Our findings suggest that RNA degradation and termination of transcription play a major role in Polycomb-mediated gene silencing in human cells.

## Results

### The rixosome associates with Polycomb

We used CRISPR/Cas9 genome editing in human embryonic kidney (HEK293FT) cells to modify the endogenous copies of *WDR18* and *NOL9* genes, which encode rixosome subunits, to express 3xFlag-WDR18 and 3xFlag-NOL9 (**Fig. 1A, Fig. S1A**). WDR18 and NOL9 are the human orthologs of the fission yeast Crb3 and Grc3 subunits of the rixosome, mutations in which disrupt heterochromatin maintenance (Shipkovenska et al., 2019). Since the rixosome plays an essential role in rRNA processing and ribosome biogenesis in nucleoli (Fromm et al., 2017), we devised a fractionation protocol to enrich for chromatin-bound rixosomes (**Fig. 1A**). We used an anti-Flag antibody to immuno-purify 3xFlag-NOL9 protein after DNA digestion with DNase I (**Fig. 1A**). Tandem Mass Tag spectrometry (TMT-MS) analysis of immunoprecipitates, identified all seven known subunits of the rixosome, including NOL9, WDR18, LAS1L, MDN1, PELP1, TEX10, and SENP3 (**Fig. 1C**). In addition, several subunits of the PRC1 (RING1B, USP7, and L3MBTL2) and PRC2 (EZH2, EED, and SUZ12) co-purified with 3xFlag-NOL9, albeit at lower efficiency than core rixosome components (**Fig. 1C**). In contrast to fission yeast (Shipkovenska et al., 2019), H3K9me-associated HP1 proteins (CBX1, 3, and 5) were not significantly enriched in the rixosome purification (**Fig. 1C**), suggesting that in human cells the rixosome may regulate H3K27me3-, rather than H3K9me3-, modified heterochromatin. Similar to Flag-NOL9 purifications, MS analysis showed that Flag-WDR18 purifications contained all seven subunits of the rixosome and, in addition, were enriched for PRC1 (RING1B, RYBP, and USP7) and PRC2 (EZH2 and RBAP46) subunits (**Fig. S1A, B**). We further validated the association of RING1B and EZH2 with NOL9 and WDR18 by immunoprecipitation and western blot analysis (**Fig. S1C, D**). Together, these results suggest that both PRC1 and PRC2 complexes physically associate with the rixosome.

**Fig. 1.**
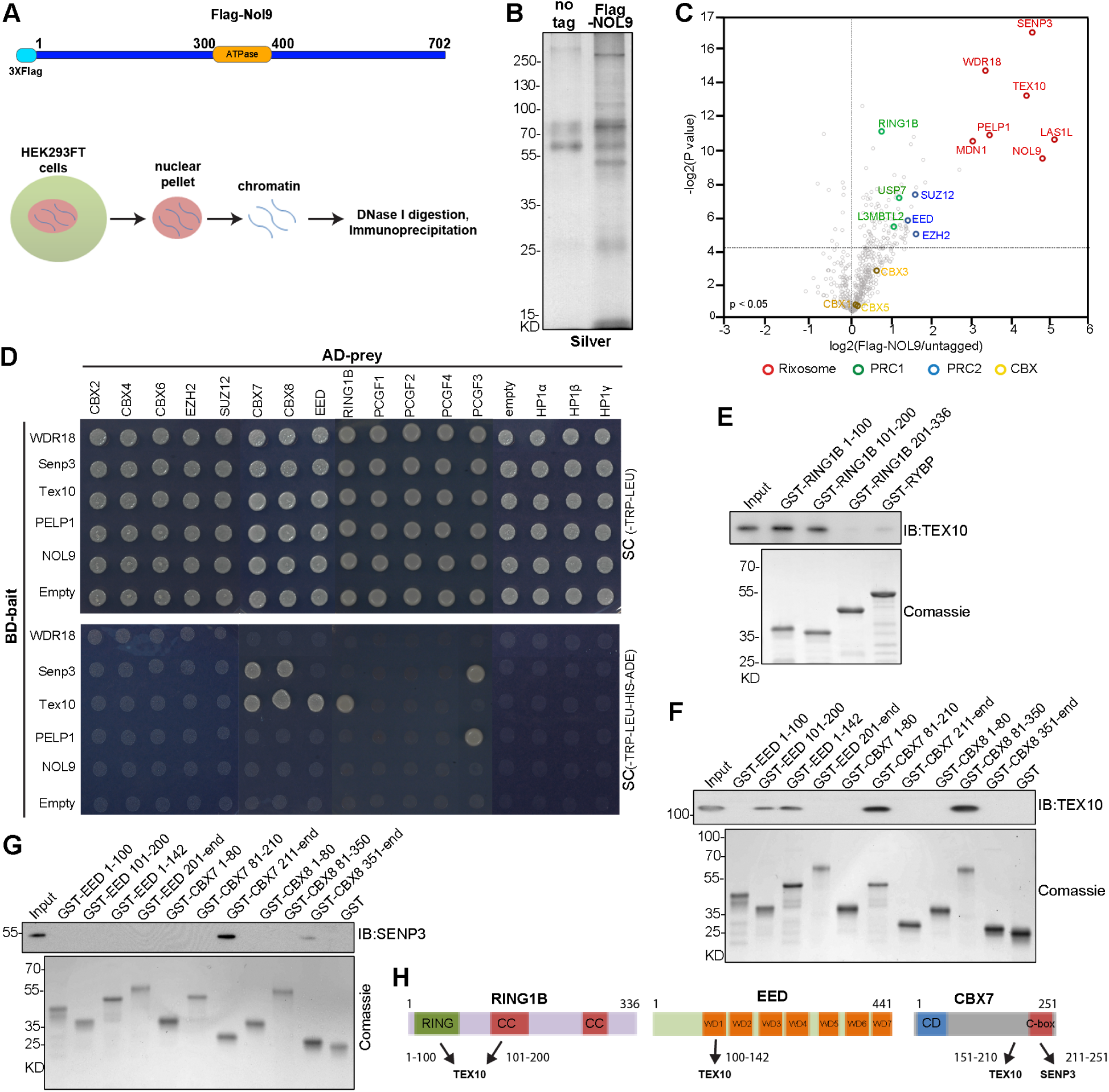
The rixosome associates with the PRC1 and PRC2 components. **(A)** Genomic tagging of *NOL9* with 3xFlag and experimental design for protein immunoprecipitation (IP) from a chromatin fraction. **(B)** Silver staining of Flag antibody immunoprecipitations from wild-type and Flag-NOL9 HEK293FT cells. **(C)** Volcano plot displaying TMT-MS results of proteins enrichment in Flag immunoprecipitations from Flag-NOL9 IP relative to untagged cells from two independent experiments. *p* value calculated by t test. The rixosome, PRC1, PCR2, and CBX proteins are highlighted. **(D)** Yeast two-hybrid assays. Yeast cells transformed with the indicated plasmids were plated onto double selective medium (Trp–, Leu–) or quadruple selective medium (Trp–, Leu–, His–, Ade–). AD, Activation Domain; BD, Binding Domain. (**E**) Pull down assays using bacterially expressed and purified TEX10 proteins incubated with beads coated with GST or GST-tagged RING1B fragments or RYBP proteins. Samples were immunoblotted (IB) with anti-TEX10 antibody. GST-tagged proteins were stained with Coomassie blue. **(F)** Pull down assays using bacterially expressed and purified TEX10 proteins incubated with beads coated with GST or indicated GST-tagged proteins. The samples were immunoblotted (IB) with anti-TEX10 antibody. GST-tagged proteins were stained with Coomassie blue. **(G)** Pull down using bacterially expressed and purified SENP3 (1-400 aa) proteins incubated with beads coated with GST or indicated GST-tagged proteins. The samples were immunoblotted (IB) with anti-SENP3 antibody. GST-tagged proteins were stained with Coomassie blue. **(H)** Summary of interactions between RING1B, EED, and CBX7 with rixosome subunits. WD, tryptophan aspartate repeat; CC, coiled-coil; CD, chromodomain; C-box, C-terminal box.

To further test interactions between Polycomb complexes and the rixosome, we carried out a series of yeast two-hybrid (Y2H) assays to identify potential direct protein-protein interactions. These assays included 13 Polycomb core subunits, 3 HP1 proteins (CBX1/3/5), and the WDR18, SENP3, TEX10, PELP1, and NOL9 subunits of the rixosome. The results indicated that SENP3 interacted with CBX7, CBX8, and PCGF3, TEX10 with RING1B, EED, CBX7, and CBX8, and PELP1 with PCGF3 (**Fig. 1D**). We observed no interactions between HP1 proteins and any of the rixosome subunits we tested (**Fig. 1D**). In support of the biochemical data (**Fig. 1B,C** and **Fig. S1A-E**), the Y2H assays demonstrate extensive interactions between the rixosome and subunits of the PRC1 and 2 complexes.

We next performed GST pull-down assays using bacterially expressed and purified proteins to reconstitute interactions between the rixosome and Polycomb complexes. We found that the N-terminal and middle regions of RING1B (amino acids 1-100 and 101-200, respectively) bound to TEX10 (**Fig. 1E, H**). Furthermore, we narrowed down the interaction of EED with TEX10 to the first WD repeat of EED (amino acids 142-200) using the GST pull-down assay (**Fig. 1F, H**) and confirmed this interaction using Y2H assays (**Fig. S1F**). We then used the Y2H and GST pull-down assays to examine the regions in CBX7 and CBX8 that bind to TEX10 and SENP3, and found that amino acids 151-210 of CBX7 and 311-350 of CBX8 bound to TEX10 (**Fig. S1G-H** and **Fig. 1G, H**), while the C-terminal domain containing the c-box domain in CBX7 and CBX8 interacted with SENP3(**Fig. S1I-J and Fig. 1H**). The rixosome therefore interacts directly with subunits of both PRC1 and PRC2.

### Nuclear and chromatin localization of the rixosome

We first used immunofluorescence staining to test for colocalization of rixosome and the EZH2 subunit of PRC2 in human embryonic stem (ES) cells. The results showed that the NOL9, WDR18, and MDN1 subunits of the rixosome localized to closely overlapping domains with EZH2 Polycomb bodies (**Fig. 2A**). To test whether the localization pattern of rixosome subunits was H3K27me3-dependent, we examined their localization patterns in ES cells carrying deletions of both the EZH1and EZH2 H3K27 methyltransferases (DKO). As shown in **Fig. 2A** and **Fig. S2A**, in the DKO ES cells, the rixosome localization pattern was altered as its subunits colocalized with the nucleolar marker NPM1. The Rixosome colocalized with EZH2 in HEK293FT and human skin fibroblast BJ-5ta cells (**Fig. S2B-C**), but not with H3K9me3 foci (**Fig. S2D**). The specificity of the antibodies used in the above experiments was verified by immuno-staining and western blot analysis after siRNA knockdowns in HEK293FT cells (**Fig. S2B, E-J**). Therefore, consistent with the biochemical and Y2H interaction data, the immuno-localization experiments support a close association between the rixosome and nuclear Polycomb bodies.

**Fig. 2.**
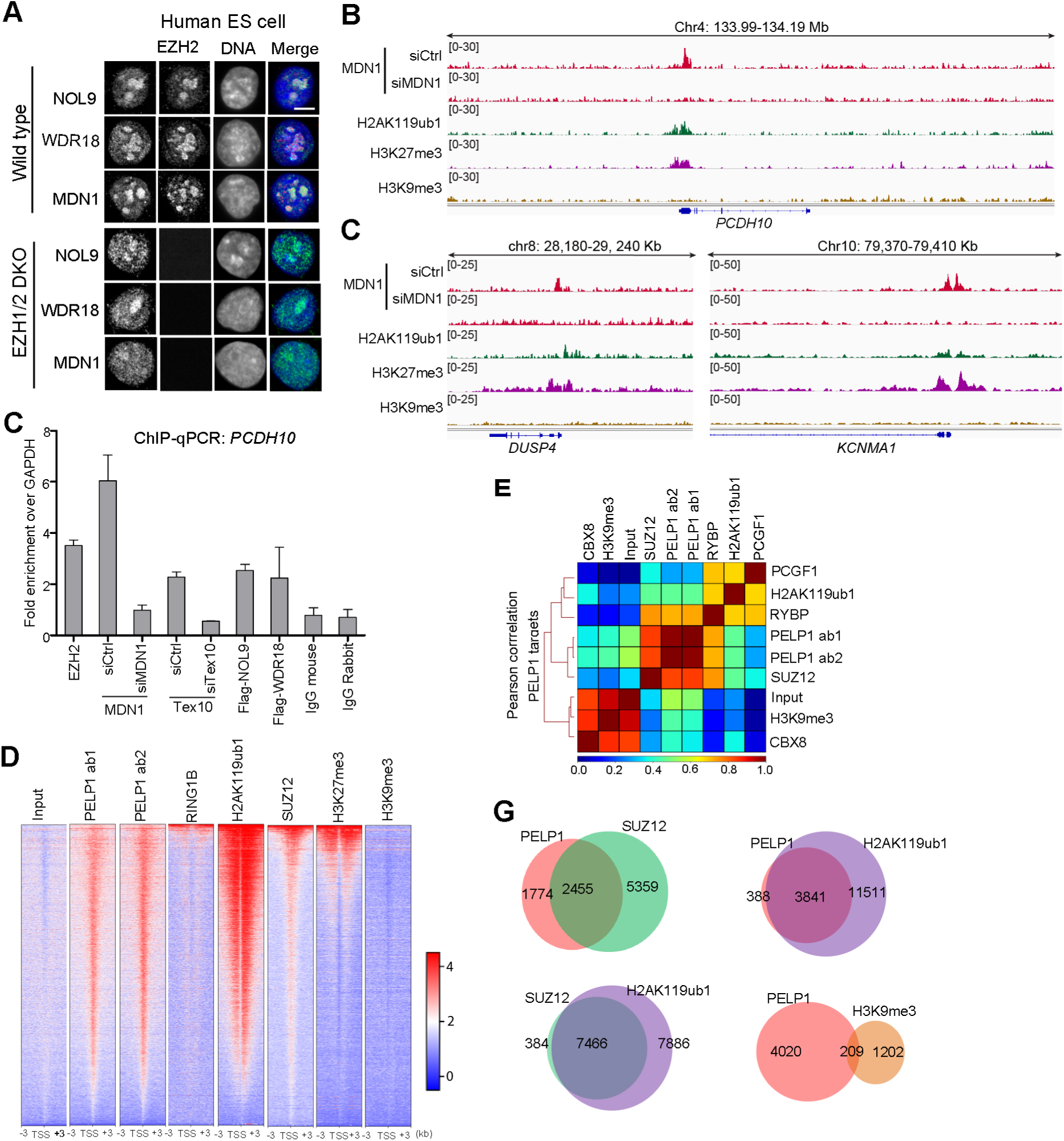
Nuclear and chromosomal colocalization of rixosome and Polycomb proteins. **(A)** Immunofluorescence of rixosome components NOL9, WDR18, and MDN1 with EZH2 in wild-type or *EZH1/2* double knockout human ES cells. DNA was stained with DAPI (blue). Scale bar, 5 μm; TPM, Transcript per Million. **(B)** Genomic snapshots of ChIP-seq reads for the MDN1 subunits of the rixosome showing colocalization with H2AK119ub1 and H3K27me3 modifications at Polycomb target genes *PCDH10, DUSP4*, and *KCNMA1* in HEK293FT cells. siRNA knockdown (siCtrl and siMDN1) was used to control for the specificity of the MDN1 antibody. **(C)** ChIP-qPCR showing localization of rixosome subunits at the *PCDH10* gene in HEK293FT cells. siRNA knockdowns were used to verify antibody specificity; mouse and rabbit IgG served as controls. Error bars represent standard deviations for three biological replicates. **(D)** Heatmaps of PELP1 (ab 1, antibody 1; ab 2, antibody 2), SUZ12, H3K27me3, H2AK119ub1, H3K9me3, and RING1B ChIP-seq in human ES cells at transcription start sites (+/-3kb) (n=41,088). **(E)** Pearson correlation of indicated ChIP-seq signal over PELP1 binding regions in human ES cells (based on peak calling, n=27, 692). **(F)** Venn diagrams showing overlap between the target genes for the indicated factors in human ES cells by ChIP-seq. For PELP1 the common PELP1 target genes (n=4229) were used for comparing.

To examine the genome-wide localization of the rixosome in human cells, we carried out chromatin immunoprecipitation followed by high throughput sequencing (ChIP-seq) in HEK293FT cells using an antibody that recognizes the MDN1 subunit of the rixosome. As a control for antibody specificity, we performed ChIP-seq on cells treated with either control (Ctrl) or MDN1 siRNAs, respectively. MDN1 specifically localized to the promoter regions of many Polycomb target genes. As examples, we observed localization of MDN1 to *PCDH10, DUSP4*, and *KCNMA1* genes, which have previously been shown to be enriched for H2AK119ub1, H3K27me3, and EZH2 (**Fig. 2B, C**). Consistent with the ChIP-seq results, ChIP followed by quantitative PCR (ChIP-qPCR) showed that the MDN1, TEX10, NOL9, and WDR18 subunits of the rixosome colocalized with EZH2 at the promoter region of the *PCDH10* Polycomb target (**Fig. 2C**). As further validation, we performed ChIP-seq in human ES cells using two different antibodies (ab1, mouse monoclonal; ab2, rabbit polyclonal) that recognize the PELP1 subunit of the rixosome. We verified the specificity of the anti-PELP1 ab2 by siRNA knockdown and immunoblotting in HEK293FT cells (**Fig. S2J**). We generated heat maps of the ChIP-seq reads on 6-kb intervals centered on transcription start sites (TSSs) and found similar enrichment patterns for PELP1, subunits of PRC1 and PRC2 (RING1B and SUZ12), and the modifications they catalyze (H2AK119ub1and H3K27me3)(**Fig. 2D**). Further, Pearson correlation analysis confirmed a close correlation between PELP1 targets and those of the PRC1 and PRC2 complexes (**Fig. 2E, Fig. S3A-B**). By contrast, the rixosome and H3K9me3, which is largely non-overlapping with H3K27me3 domains in human cells, did not show similar enrichment patterns (**Fig. 2D, E**). Consistent with these results, we observed a similar enrichment pattern for the MDN1 subunit of the rixosome and H3K27me3 and H2AK119ub1 modifications (**Fig. S3C**).

At the genome-wide level (+/-2 kb from TSS), PELP1 ab1 and ab2 ChIP-seq peaks (4229) showed good overlap (∼70%, 4229 of 5910)(**Fig. S3D**). Comparison of these targets with Polycomb targets showed that 58% and 91% of PELP1 targets were also bound by SUZ12 and H2AK119ub1, respectively (**Fig. 2G**). There was similarly good overlap between PELP1 and MDN1 targets and H3K27me3 and H2AK119ub1 targets, but not with H3K9me3, in both ES and HEK293FT cells (**Fig. S3 E-G**). The rixosome therefore colocalizes with the Polycomb machinery to target genes, raising the possibility that it contributes to controlling their expression.

### Rixosome in Polycomb-mediated silencing

We next investigated whether the rixosome has a direct role in silencing of Polycomb target genes. The rixosome plays critical roles in rRNA processing (rRNA), and as expected, we failed to obtain *NOL9* and *WDR18* knockout human ES cell lines. We therefore used transient siRNA knockdown of rixosome subunits at timepoints that do not affect growth and proliferation to study the role of the rixosome in regulation of transcription. We successfully knocked down rixosome subunits (NOL9, LAS1L, WDR18, MDN1, and SENP3) and EZH2 (**Fig. S2E-J, Fig. S4A**). Growth curve of cells after knockdown of LAS1L, NOL9, and EZH2 showed that 48 h of siRNA treatment did not affect cell proliferation (**Fig. S4B**). Using RT-qPCR, we found that 48 h knockdown of NOL9, LAS1L, WDR18, MDN1, and SENP3 elevated the expression of PCDH10 by ∼ 3 fold, similar to the level of depression observed in EZH2 knockdown cells, while knockdown of SUV39H1 had no effect (**Fig. 3A**), suggesting that the rixosome was required for silencing of this Polycomb target gene.

**Fig. 3.**
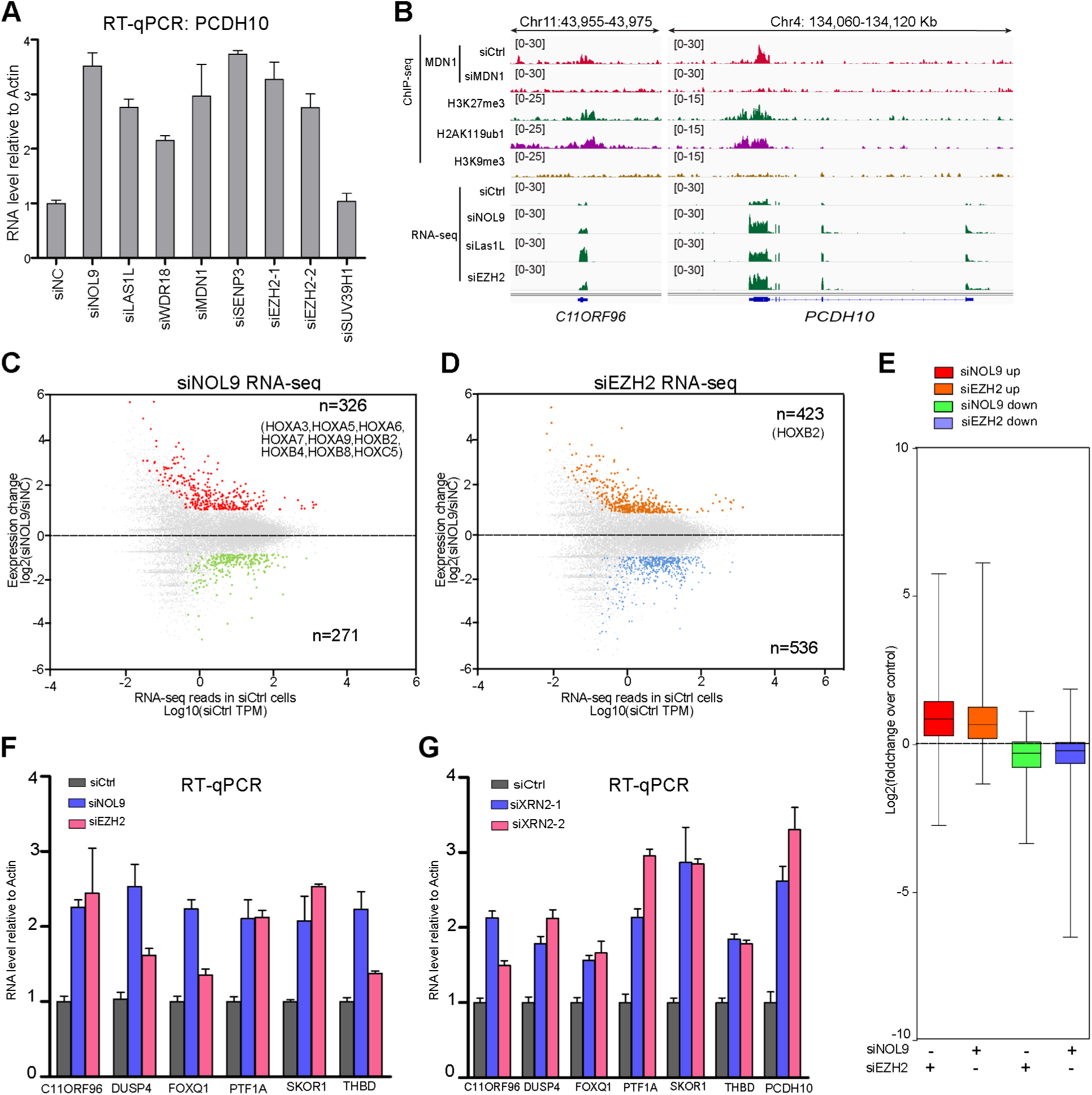
The rixosome is required for repression of Polycomb target genes. **(A)** RT-qPCR assays showing that siRNA knock down of rixosome subunits results in increased expression of *PCDH10* in HEK293FT cells. *ACTB* encoding actin served for normalization. Error bars represent standard deviations for three biological replicates. **(B)** Genomic snapshots of ChIP-seq and RNA-seq reads showing that the effect of siRNA knockdown of NOL9, LAS1L, and EZH2 on the expression of Polycomb target genes *C11ORF96* and *PCDH10* in HEK293FT cells. **(C)** Dot plot of log2-fold changes in gene expression (RNA-seq) in NOL9 knockdown cells.m Significant gene expression changes (p-adj < 0.05, fold change > 2.0) are shown in red (upregulated) and green (downregulated). The upregulated *HOX* genes are indicated. **(D)** Dot plot of log2-fold changes in gene expression (RNA-seq) in EZH2 knockdown cells. Significant gene expression changes (p-adj < 0.05, fold change > 2.0) are shown in brown (upregulated) and blue (downregulated). The upregulated *HOX* gene is indicated. **(E)** A boxplot show log2-fold changes over control (RNA-seq) of indicated gene sets with indicated siRNA treatment. upregulated (Up), downregulated (Down). **(F)** RT-qPCR of Polycomb target genes *FOXQ1, DUSP4, PTF1A, C11ORF96, THBD, and SKOR1* in NOL9 or EZH2 siRNA knockdown cells. RNA expression levels were normalized to housekeeping gene *ACTB*. Error bars represent standard deviations for three biological replicates. **(G)** RT-qPCR showing that knockdown of XRN2 with 2 different siRNAs results in increased expression of the indicated Polycomb target genes. RNA expression levels were normalized to housekeeping gene *ACTB*. Error bars represent standard deviations for three biological replicates.

To assess the global effects of rixosome knockdowns on gene expression, we carried out poly(A)-selected RNA-seq. As shown in **Fig. 3B, C** and **Fig. S4C, D**, knockdown of NOL9 and LAS1L resulted in increased expression of Polycomb target genes. We further showed that the effect of siNOL9 on upregulation of several target genes was rescued by the reintroduction of siRNA-resistant wild-type NOL9 (HA-NOL9 WT) but not a mutant catalytically dead NOL9 (HA-NOL9 K312A)(**Fig. S4E-F**). The polynucleotide kinase activity of NOL9 is therefore required for its silencing function. Globally, at a cutoff fold change of >2 and false discovery rate (FDR) < 0.05, a 2-day knockdown of NOL9 derepressed 326 genes and repressed 271 genes, while knockdown of EZH2 resulted in derepression of 423 genes and repression of 536 genes (**Fig. 3C, D**). Although 113 genes were commonly derepressed in both NOL9 and EZH2 KD (35% overlap)(**Fig. S5A-C**), a larger number of HOX genes were derepressed in NOL9 knockdown cells (9) than in EZH2 knockdown cells (1), suggesting that the rixosome may play a more critical role in HOX gene silencing than EZH2 in HEK293FT cells (**Fig. 3C, D**). Consistent with co-regulation of target genes by rixosome and Polycomb, boxplot analysis showed that siEZH2 knockdown (KD) upregulated the genes that were also upregulated by siNOL9 KD, while genes that were downregulated by siNOL9 were not affected by knockdown of EZH2 (**Fig. 3E**). Enrichment assays of NOL9 KD derepressed gene set (326 genes) and EZH2 KD depressed gene set (423 genes) in EZH2 KD RNA-seq data and NOL9 KD RNA-seq data respectively further support the positive correlation of gene expression change between NOL9 and EZH2 KD (**Fig. S5D and E**). Furthermore, the gene expression levels of NOL9 KD show positive correlation with EZH2 KD but not control siRNA treatment (**Fig. S5F and G**). We also performed gene ontology analysis of genes that were upregulated in NOL9 and EZH2 knockdown cells and found similar sets of genes involved in development, cell differentiation, and embryo morphogenesis were upregulated with each knockdown (**Fig. S6A-B**). Together, these results demonstrate that the rixosome and Polycomb complexes repress a common set of genes in HEK293FT cells.

In fission yeast, the rixosome promotes the degradation of target RNAs by the conserved 5’-3’ exoribonuclease Dhp1, the homolog of the human XRN2 protein (Shipkovenska et al., 2019). To test whether the rixosome and XRN2 act in the same pathway in human cells, we knocked down XRN2 in human HEK293FT cells and found that several targets of the rixosome and Polycomb pathways were expressed at elevated levels upon knockdown of XRN2 with 2 different siRNAs (**Fig. 3F, G** and **Fig. S7**). The rixosome is therefore likely to promote XRN2-mediated degradation of its targets in human cells.

### Rixosome repression of HOX genes

The *HOX* genes encode conserved master transcription regulators that specify positional information along the body axis and control cell type-specific gene expression, and are amongst the best-studied targets of Polycomb complexes (Montavon and Duboule, 2013). We therefore investigated the extent to which the rixosome regulates *HOX* gene expression. The human genome contains 39 *HOX* genes located at four distinct genomic cluster (Montavon and Duboule, 2013, Duboule, 2007). We quantified the expression of all *HOX* genes, except *HOXB1*, using our RNA-seq data. Surprisingly, boxplot analysis and examination of individual *HOX* gene RNAs showed that the *HOX* genes were upregulated to a higher extent with NOL9 and LAS1L knockdown than with EZH2 knockdown in HEK293FT cells (**Fig. 4A, B** and **Fig. S8A**). This observation was consistent with ChIP data, which showed that *HOXA* genes in HEK293FT cells were not enriched for H3K27me3, but instead were marked by PRC1-catalyzed H2AK119ub1 (**Fig. 4B, C**). Moreover, the expression of the *HOX* genes was elevated by knockdown of the E3 ubiquitin ligase, RING1B, as well as knockdowns of rixosome subunits and XRN2, but not EZH2, suggesting that they were repressed by the PRC1/H2AK11119ub1 system and the rixosome, despite lacking H3K27me3 (**Fig. S7B, C**). To test whether the rixosome localization to HOX genes required H2AK119ub1, we used ChIP-qPCR to analyze the localization of the MDN1, PELP1, and TEXT10 subunits of the rixosome in RING1B knockdown cells. The results showed that depletion of RING1B, but not the EED subunit of PRC2, reduced the binding of rixosome subunits to the *HOX* genes, indicating that PRC1-mediated H2AK119ub1 was required for targeting the rixosome to HOX genes (**Fig. 4C**). Consistent with the paucity of H3K27me3 at *HOX* genes, ChIP-qPCR showed that H2AK119ub1 at these loci was not affected by knockout of EED, which is required for PRC2-mediated H3K27 methylation (**Fig. 4D**)(Margueron et al., 2009). In contrast to the *HOX* genes, deletion of PRC2 subunits significantly reduced Rixosome enrichment at loci that were highly enriched for PRC2 and H3K27me3 (**Fig. S10A**). We conclude that at some loci PRC1-mediated H2AK119ub1 is sufficient for recruitment of the rixosome, while at other loci both PRC1 and PRC2 play roles in chromatin targeting the rixosome to chromatin. Finally, the rixosome was associated with *HOX* genes in four other cell types, including human ES cell, human neural progenitor cells, mouse C2C12, and human HeLa cells, demonstrating that its localization to Polycomb target genes was not cell-type specific (**Fig. S9A-D**).

**Fig. 4.**
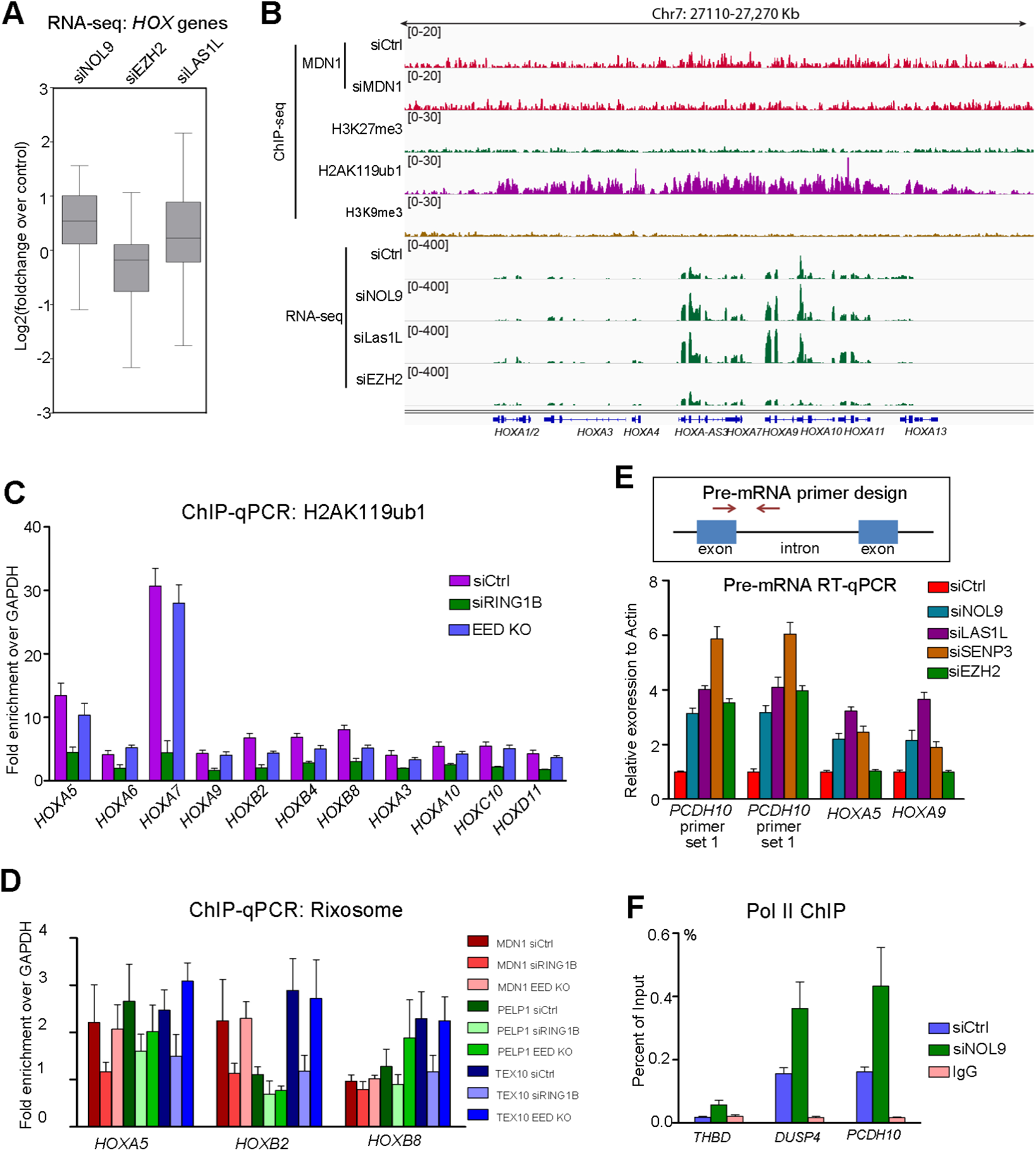
Rixosome represses HOX genes. **(A)** Boxplot analysis of RNA-seq results with the indicated siRNA treatment in HEK293FT cells showing relative changes in the expression of 37 *HOX* genes (all, except *HOXB1*, TPM=0 in siCtrl cells). **(B)** Genomic snapshot showing increased expression of genes in the *HOXA* locus upon siRNA knockdown of rixosome subunits NOL9 and LAS1L, but not EZH2, showing in HEK293FT cells. ChIP-seq reads for MDN1 (siCtrl and siMDN1), H3K27me3, H2AK119ub1, and H3K9me3 are presented for reference. **(C)** ChIP-qPCR analysis for H2K119ub1 at indicated *HOX* genes in siCtrl, siRING1B, or EED KO HEK293FT cells. Data represent three biological replicates. **(D)** ChIP-qPCR analysis of MDN1, PELP1, and TEX10 at indicated HOX genes in siCtrl, siRING1B, or EED knockout (KO) HEK293FT cells. Error bars represent standard deviations for three biological replicates. **(E)** Top, schematics of primer design for RT-qPCR quantification of pre-mRNA levels. Bottom, RT-qPCR results showing the effect of specific siRNA knockdowns on pre-mRNA levels for the indicated Polycomb-rixosome target genes in HEK293FT cells. Pre-mRNA expression levels were normalized to the housekeeping gene *ACTB*. Error bars represent standard deviations for three biological replicates. **(F)** ChIP-qPCR results showing that NOL9 knockdown results in increased RNA polymerase II (Pol II) at the indicated Polycomb-rixosome target genes in HEK293FT cells. Control siRNA knockdown (siCtrl) and mouse IgG served as controls. Error bars represent standard deviations for three biological replicates.

Localization of the rixosome to Polycomb target genes and its requirement for Polycomb-mediated repression suggest that it may act co-transcriptionally to degrade nascent transcripts to promote RNA polymerase II (Pol II) termination within these domains. To test this hypothesis, we asked whether pre-mRNA and RNA polymerase II (Pol II) accumulated at target genes upon rixosome and Polycomb knockdowns. As shown in **Fig. 4E**, knockdown of the NOL9, LAS1L, and SENP3 subunits of the rixosome resulted in accumulation of pre-mRNA at the *PCDH10, HOXA5*, and *HOXA10* genes. By contrast, knockdown of EZH2 alone increased pre-mRNA accumulation at the *PCDH10* gene, which is associated with both H2AK119ub1 and H3K27me3, but not at the *HOXA5* and *HOXA10* genes, which are solely associated with H2AK119ub1 (**Fig. 4E**). Furthermore, knockdown of NOL9 increased Pol II occupancy at promoters of Polycomb target genes, *THBD, DUSP4*, and *PCDH10* (**Fig. 4F**). Importantly, 48 h depletion of the rixosome did not affect H2AK119ub1 or H3K27me3 levels at target genes, suggesting that the rixosome regulates Pol II termination and release at a step downstream of the above histone modifications (**Fig. S10A-D**).

## Discussion

The principal mechanism of silencing by Polycomb complexes is thought to involve chromatin compaction (Schuettengruber et al., 2017, Simon and Kingston, 2013). In both flies and mammals, subunits of the PRC1 complex can condense nucleosomal arrays *in vitro* (Francis et al., 2004, Shao et al., 1999, Grau et al., 2011) and, in mammals, PRC2 alone has *in vitro* chromatin compaction activity (Margueron et al., 2008, Terranova et al., 2008). Moreover, recent studies show that the CBX2 subunit of the PRC1 complex, which mediates its chromatin compaction activity, also promotes liquid-liquid phase separation (LLPS) both *in vitro* and *in vivo* (Plys et al., 2019, Tatavosian et al., 2019). Our identification of a role for the rixosome in silencing of Polycomb target genes suggests that RNA degradation plays an important role in the mechanism of Polycomb-mediated gene silencing. In our experiments, knockdown of rixosome subunits does not seem to disrupt Polycomb bodies (**Fig. S2**) or reduce the levels of Polycomb-mediated histone modifications (**Fig. S8H-J**), leading us to propose that rixosome-mediated RNA degradation and transcription termination are critical at a step downstream of histone modification and condensate formation (**Fig. 5**).

**Fig.ure 5.**
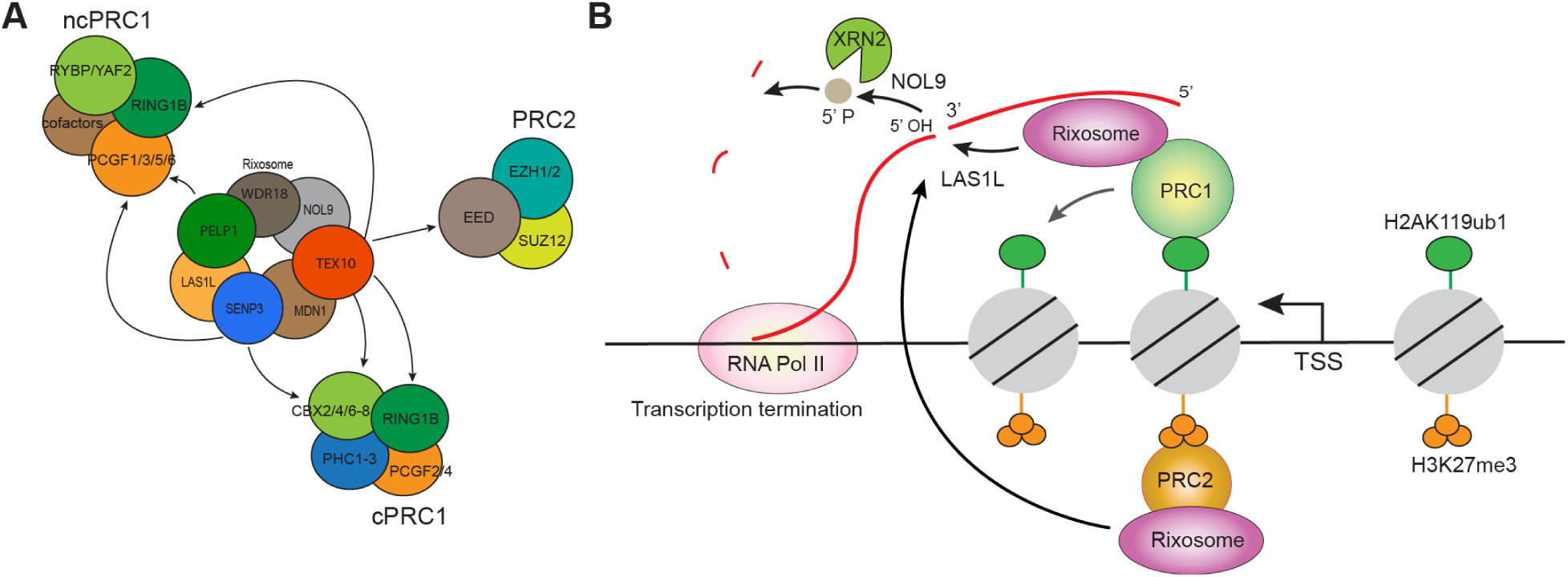
Summary of rixosome interactions with Polycomb complexes and model for rixosome-dependent degradation of nascent RNA and transcription termination. (**A**) The rixosome interacts with the canonical and non-canonical PRC1 complexes (cPRC1 and ncPRC1), and with PRC2. (**B**) The rixosome is recruited by Polycomb complexes to mediate RNA degradation and transcription termination.

The rixosome also mediates heterochromatic RNA degradation within H3K9me domains in fission yeast (Shipkovenska et al., 2019). Its association with H3K27me3, rather H3K9me3, domains in human cells, suggests that the complex can be recruited to different types of repressed chromatin via exchangeable protein-protein interaction surfaces. The conservation of RNA degradation in silencing functions of different types of heterochromatin domains suggests that RNA degradation plays a highly conserved and critical function in heterochromatin-mediated gene silencing. In this regard, another conserved RNA processing and degradation machinery that localizes to H3K9me heterochromatin involves nuclear RNAi (Holoch and Moazed, 2015). The roles of RNAi extend beyond RNA degradation and include amplification mechanisms that provide a memory of the initial trigger that induces RNAi (Yu et al., 2018, Kowalik et al., 2015). In addition, in contrast to the rixosome, which appears to act solely downstream of repressive chromatin modifications, nuclear RNAi feeds back on the chromosome to promote and reinforce H3K9me and heterochromatin formation, making it difficult to parse its degradation-dependent silencing functions from its roles in memory and establishment of heterochromatin. Our findings raise the possibility that nuclear RNAi first evolved to degrade and silence nascent transcripts, via direct localization to nascent chromatin-bound transcripts (Motamedi et al., 2004, Buhler et al., 2006), and only later gained the capability to also induce heterochromatin formation. We further propose that RNA degradation, associated with the rixosome, RNAi, or other pathways, is an ancient mechanism of heterochromatin-mediated gene silencing, which may have preceded the evolution of mechanisms that lead to compaction, condensate formation, and transcriptional gene silencing.

## Materials and Methods

### Methods

#### Plasmid construction

Rixosome subunits (NOL9, WDR18, PELP1, TEX10, WDR18,), PRC2 subunits (EZH2, EED, SUZ12), PRC1 subunits (RING1B, PCGF1-4) and CBX1-8 cDNAs were amplified from human ES cell cDNA library and inserted to pGAD-T7 (Takara, 630442) and pGBK-T7 (Takara, 630443) plasmids for yeast two-hybrid assays. NOL9 siRNA resistant cDNA was generated by PCR. The siRNA target sequence was mutated from 5’-AGACCTAAGTTCTGTCGAA-3’ to 5’-CGGCCGAAATTTTGCAGGA-3’ and integrated into the pCI (Promega, E1731) plasmid for ectopic protein expression. For bacteria protein expression, cDNA was integrated to pGEX-6P-1 (GE Healthcare, 28-9546-48).

#### Cell culture

C2C12 (ATCC, CRL-1772), HeLa (ATCC, CCL-2), and HEK293FT (ThermoFisher, R70007) cells were cultured in DMEM containing 10% fetal calf serum. BJ-5ta (ATCC CRL-4001) were cultured in a mixed medium (80% DMEM and 20% Medium 199) containing 10% fetal calf serum. Human embryonic stem cells were cultured as previously described (Chen et al., 2011). Briefly, cells cultures on 0.08 mg/ml matrigel coated plates with DMEM/F12 (containing 5 μg/ml insulin and 10 μg/ml, 0.1 μg/ml FGF2, 1.7 ng/ml TGFβ1, 10 μg/ml transferring). Neural progenitor cells were derived from human ES cells by NGN2 induction. The donor plasmid pAAVS1-TRE3G-NGN2 was constructed by replacing *EGFP* with human *NGN2* in plasmid AAVS1-TRE3G-EGFP (Addgene plasmid # 52343). The donor plasmid pAAVS1-TRE3G-NGN2 (5 μg), hCas9 (Addgene plasmid # 41815) (2 μg), and gRNA_AAVS1-T2 (Addgene plasmid # 41818) (2 μg) were electroporated into 1×10^6^ human ES H9 cells. The cells were treated with 0.25 μg/ml Puromycin for 7 days and colonies were genotyped. The primers for 5’ junction PCR were 5’-CTCTAACGCTGCCGTCTCTC and 5’-TGGGCTTGTACTCGGTCATC. The primers for 3’ junction PCR were 5’-CACACAACATACGAGCCGGA and 5’-ACCCCGAAGAGTGAGTTTGC. The primers for locus were PCR: 5’-AACCCCAAAGTACCCCGTCT and 5’-CCAGGATCAGTGAAACGCAC. For human ES cell induction to neural progenitor cells, cells were plated at 2×104/cm2 on Matrigel-coated plates in DMEM/F12 supplemented with 1x N2, 1x NEAA (ThermoFisher Scientific), human Brain-derived neurotrophic factor (BDNF, 10 ng/ml, PeproTech), human Neurotrophin-3 (NT-3, 10 ng/l, PeproTech), mouse laminin (0.2 μg/ml, Cultrex), Y-27632 (10 μM, PeproTech) and Doxycycline (2 μg/ml, Alfa Aesar) on Day 0. On Day 1, Y-27632 was removed. Starting on Day 2, Neurobasal medium was used, which was supplemented with 1x B27 and 1x Glutamax (ThermoFisher Scientific) containing BDNF, NT-3 and 1 μg/ml Doxycycline. Starting on Day 4, half of the medium was replaced every other day. On day 6, the cells were harvested for downstream assays.

#### RNAi

For siRNA-mediated knockdown, Lipofectamine™ 3000 (Invitrogen) and siRNA (200 nM) were used to transfect the cells by following the manufacturer’s instructions. All the siRNAs were synthesized by Dharmacon and are listed in Table S2.

#### CRISPR/Cas9-mediated human genome editing

Small guide RNA was synthesized via in vitro transcription by using MAXIscript T7 transcription kit (ThermoFisher, AM1312). CRISPR/Cas9 protein was purified and provided by Initiative for Genome Editing and Neurodegeneration Core in the Department of Cell Biology at Harvard Medical School. Oligo DNA template (synthesized by IDT), guide RNA, and CRISPR/Cas9 protein were delivered to cells by electroporation with Neon transfection system (ThermoFisher). Clones were screened by PCR and Miseq sequencing (Illumina).

#### Immunofluorescence

Cells were placed on plates with cover slides. For human ES cell staining, slides were coated with Matrigel before plating. ES cells were digested with Accutase (STEMCELL Technologies, 07922) to separate cells. Rock kinase inhibitor Y-27632 (10 μM, PeproTech) was used to promote cell attachment on day 0. Y-27632 was removed on day 1. On day 2, the ES Cells were used for immunostaining. Cells were first washed with PBS, and fixed and permeabilized with methanol for 8 min at −20°C. Cells were then incubated for 4-10 h at 4°C with primary antibodies in PBS containing 4% bovine serum, which was followed by staining with secondary antibodies and 1 μg/ml DAPI. A confocal microscope (Nikon, Ti w/perfect focus&spinning disk) equipped with a 60×/1.40 NA objective lens was used to image cells. Images were post-processed with Image J (NIH) and photoshop (Adobe) software.

#### Immunoprecipitation and Mass spectrometry Analysis

To prepare chromatin-enriched fractions, cells were washed with PBS and then resuspended in ice-cold hypotonic buffer (10 mM HEPES, pH7.9, 1.5 mM MgCl_2_, 10 Mm KCl, 0.2 mM PMSF, 0.2 mM DTT) and incubated on ice for 10 min. Cell membranes were then disrupted by douncing for 10 times. Nuclei were pelleted by centrifugation at 2,000xg for 10 min, resuspended in cell lysis buffer (50 mM Hepes, pH 7.4, 150 mM NaCl, 1 mM MgCl_2_, 1 mM EGTA, and 0.5% Triton X-100) by pipetting for 3 min, and pelleted by centrifugation at 2,000xg for 10 min to obtain a chromatin fraction. The chromatin pellet was resuspended in IP buffer (50 mM Hepes, pH 7.4, 250 mM NaCl, 10% Glycerol, 1 mM MgCl_2_, 1 mM EGTA, and 1% Triton X-100) containing protease inhibitor cocktail (5056489001, Sigma) and 1 mM DNase I. Chromatin was digested for 2 h at 4°C and centrifuged at 10,000xg for 10 min. The supernatant was then incubated with specific antibodies and immune complexes were collected using Dynabeads Protein A/G (ThermoFisher). For silver staining, samples were run on a 5%-20% Bis-Tris SDS-PAGE gel (BioRad) and stained with SilverQuest Silver Staining kit (Invitrogen) according to the manufacturer’s instructions. For immunoblotting, beads were boiled for 5 min in SDS loading buffer. For mass spectrometry analysis, proteins were eluted with 0.5 M NH_4_OH and dried to completion in a speed vac.

Dried protein samples were digested in 200 mM EPPS buffer pH 8.5 with trypsin (Promega #V5111). Digests contained 2% acetonitrile (v/v) and were performed at 37°C overnight. Digests were labeled directly with TMT10 plex reagents (ThermoFisher Scientific, #90406). Labeling efficiency was checked by mass spectrometry. After hydroxylamine-quenching (0.3% v/v) for 15 minutes, reactions were mixed and acidified and solvent evaporated to near completion by speed vac. Samples were then fractionated by alkaline reversed phase chromatography (ThermoFisher #84868) into 12 fractions eluted with 10%, 12.5%, 15%, 17.5%, 20%, 25%, 30%, 35%, 40%, 50%, 65% and 80% acetonitrile. Fractions were pooled into 6 fractions (1+7, 2+8, 3+9, 4+10, 5+11, 6+12), dried down, stage-tipped and analyzed by mass spectrometry on an Orbitrap Lumos instrument (Thermo Scientific).

Relative quantification followed a multi-notch SPS-MS^3^ method. Prior to injection, peptides were separated by HPLC with an Easy-nLC 1200 liquid chromatography system using 100 μm inner diameter capillaries and a C_18_ matrix (2.6 μM Accucore C_18_ matrix, ThermoFisher Scientific). Peptides were separated with 4-hour acidic acetonitrile gradients. MS^1^ scans were measured by orbitrap recording (resolution 120,000, mass range 400-1400 Th). After CID (collision induced dissociation, 35%), MS^2^ spectra were collected by iontrap mass analyzer. After SPS (synchronous precursor selection), TMT reporter ions were generated by HCD (high-energy collision-induced dissociation, 55%) and quantified by orbitrap MS^3^ scan (resolution 50,000 at 200 Th).

Spectra were searched with an in-house written software based on Sequest (v.28, rev. 12) against a forward and reversed human proteome database (Uniprot 07/2014). Mass tolerance for searches was 50ppm for precursors and 0.9 Da for fragment ions. Two missed tryptic cleavages per peptide were allowed and oxidized methionine (+15.9949 Da) was searched dynamically. For a peptide FDR (false discovery rate) of 1%, a decoy database strategy and linear discriminant analysis (LDA) were applied. The FDR for collapsed proteins was 1%. Proteins were quantified by summed peptide TMT s/n (signal/noise) with a sum s/n >200 and an isolation specificity of >70%. Details of the TMT workflow and sample preparation procedures were described recently (Navarrete-Perea et al., 2018).

#### GST Pulldown and Immunoblotting

Proteins for GST Pulldown assays were expressed in BL21 Codon Plus E. coli (Agilent Technologies) with 200 μM IPTG induction at 16 °C overnight. Bacteria were then harvested and washed with cold PBS, and sonicated (Branson sonicator) for 1 min with 20% amplitude at 4°C. Sonicated samples were centrifuged at 20,000 xg for 10 min, and the supernatant was added to 0.5 mL Glutathione Sepharose 4B resin (GE Healthcare, 17075605), which was equilibrated with PBS. GST-tagged proteins were incubated with the resin for 2 h at 4°C. The resin was then washed 6 times with PBS containing 1% Triton 100. To remove the GST tag, bead-coupled proteins were incubated with PreScission Protease (GE Healthcare, 27-0843-01) in reaction buffer (50 mM Tris-HCl, Ph7.0, 150 mM NaCl, 1 mM EDTA, 1 mM DTT) for 2 h at 4°C. The GST-tagged PreScission Protease was removed using Glutathione Sepharose 4B resin.

For GST Pulldown assays, 10 μl 50% slurry of Glutathione Sepharose 4B was used for each sample. GST or GST-tagged proteins (0.1 μM) were incubated with untagged proteins (0.1 μM) in 1 ml PBS (137 mM NaCl, 2.7 mM KCl, 8 mM Na2HPO4, and 2 mM KH2PO4, Ph7.4) containing 0.5% Triton 100 overnight at 4°C. Beads were washed 3 times with PBS containing 0.5% Triton 100, resuspended in SDS protein buffer, and boiled for 5 min. Input (2-5%) and bound proteins (10-50%) were run on 4-20% gradient SDS-PAGE gel. SDS-PAGE was performed to separate proteins for 2 h at 80 Volts, and proteins were transferred to a PVDF membrane (Millipore). The membranes were blocked in 3% milk in PBS with 0.2% Tween-20, and sequentially incubated with primary antibodies and HRP-conjugated secondary antibodies, or directly incubated with HRP-conjugated primary antibodies for chemiluminescence detection.

#### RT-qPCR

Total RNA was extracted using the RNeasy Plus kit (74134, Qiagen) and reverse transcribed into cDNA using gene-specific primers and reverse transcription kit (18090010, ThermoFisher). cDNA was analyzed by running PCR on a QuantStudio 7 Flex Real Time PCR System (Applied Biosystem). All reactions were performed using 10 ng RNA in a final volume of 10 μl. PCR parameters were 95°C for 2 min and 40 cycles of 95°C for 15s, 60°C for 15s, and 72°C for 15 s, followed by 72°C for 1 min. All the qPCR data presented were at least three biological replicates. Primer sequences are presented in Table S1.

#### Chromatin immunoprecipitation (ChIP)

ChIP was performed as previously described with minor modifications (Nowak et al., 2005). Cells for ChIP were cultured in 15 cm plates. Cell were first washed with cold PBS, crosslinked at room temperature with 10 mM DMP (ThermoFisher Scientific) for 30 min, and then 1% formaldehyde (ThermoFisher Scientific) for 15 min. Crosslinking reactions were quenched by addition of 125 mM glycine for 5 min. Crosslinked cells were separated by 3 min treatment of 0.05% trypsin (Gibco), and then washed with cold PBS 3 times. In every wash, cells were centrifuged for 3 min at 1000xg at 4°C. Cell were then resuspended in sonication buffer (pH 7.9, 50 mM Hepes, 140 mM NaCl, 1 mM EDTA, 1% Triton, 0.1% Sodium deoxycholate, and 0.5% SDS) and sonicated to shear chromatin into ∼300 bp fragments using a Branson sonicator. Sonicated samples were diluted 5-fold with ChIP dilution buffer (pH 7.9, 50 mM Hepes, 140 mM NaCl, 1 mM EDTA, 1% Triton, 0.1% Sodium deoxycholate) to obtain a final concentration of 0.1% SDS. Diluted samples were centrifuged at 13,000 rpm for 10 min. The supernatant was pre-cleared with protein A/G or Dynabeads M-280 Streptavidin beads (ThermoFisher) and immunoprecipitated for 3-12 h using 3 μg antibodies and 40 μl protein A/G or Dynabeads M-280 Streptavidin beads. The beads were washed twice with high salt wash buffer A (pH 7.9, 50 mM Hepes, 500 mM NaCl, 1 mM EDTA, 1% Triton, 0.1% Sodium deoxycholate, and 0.1% SDS), and once with wash buffer B (pH 7.9, 50 mM Hepes, 250 mM LiCl, 1 mM EDTA, 1% Triton, 0.1% Sodium deoxycholate, 0.5% NP-40). The bound chromatin fragments were eluted with elution buffer (pH 8.0, 50 Mm Tris, 10 mM EDTA, 1% SDS) for 5 min at 65°C. Eluted DNA-proteins complexes were treated with RNase A and crosslinks were reversed overnight at 65°C. Proteinase K was then added to digest proteins for 1 h at 55°C. DNA was further purified using PCR Purification Kit (QIAGEN) and analyzed by PCR on a QuantStudio 7 Flex Real Time PCR System (Applied Biosystem). PCR parameters were 95°Cfor 2 min and 40 cycles of 95°C for 15s, 60°C for 15s, and 72°C for 15 s, followed by 72°C for 1 min. All the ChIP-qPCR data presented were at least three biological replicates. Primer sequences are in the table. Error bar represent standard deviation (three biological replicates).

For ChIP-seq, sequencing library was constructed using TruSeq DNA sample Prep Kits (Illumina) and adapter dimers were removed by agarose gels electrophoresis. Sized selected and purified DNA libraries were sequenced on an Illumina Hiseq 2500 machine (Bauer core facility at Harvard University) to obtain 50 bp single-end reads. ChIP-seq reads were quality controlled with fastqc (v0.11.5) and mapped to the human genome reference (GRCh37/hg19) using bowtie2 (v2.2.9) with default parameters. All ChIP-seq peaks were called with macs2 (v2.1.1.20160309) and broad peak calling option using corresponding input files for background normalization. For MDN1 ChIP in human embryonic kidney 293 and PELP1 ChIP in human ES cells, peaks identified in two different biological replicates were used for further analysis. Peak calling results were visualized with the IGV browser. To analyze read density at TSS regions, we made heatmap and metaplot of ChIP-seq samples. TSS was centered in the regions plotted and data were tabulated with the same distance relative to TSS. Matrix files were generated using computematrix function of deeptool (v/3.0.2). Based on generated matrix file, heatmaps were generated by PlotHeatmap function, and profiles were generated by plotprofile function.

To analyze read density and correlation between different ChIP-seq samples, we performed Pearson correlation analysis. Reads density was analyzed at PRC-bound sites with multiBigwigSummary function from deeptool (v/3.0.2) to get a npz matrix file. The heatmap of Pearson correlation is generated by plotCorrelation function of deeptool (v/3.0.2). To identify target genes in each ChIP-seq library, intersect function of Bedtool (2.27.1) was used to analyze peak files. Genes containing peaks at promoter regions were considered as targets. Promoter region was defined as +/-2k from TSS. Overlapped target gene lists were also generated by bedtools intersect function. Venn diagrams were made based on the number of overlapped target gene.

The sources of published ChIP-seq data used in this study are listed in Table S4.

#### RNA-seq

Total RNA was isolated from human cells with an RNA purification kit (Qiagen, 74134) and genomic DNA was removed by genomic DNA binding columns in the kit. Two μg of total RNA was used to start RNA-seq library construction. Poly(A)-containing mRNA was isolated by poly-A selection beads and further reverse transcribed to cDNA. The resulting cDNA was ligated with adapters, amplified by PCR, and further cleaned to obtain the final library. Libraries were sequenced on an Illumina Hiseq machine (Novogene) to obtain 150 bp paired-end reads. RNA-seq reads were mapped to the human genome (GRCh37/hg19) with Kallisto (version 1.15.0). Transcript abundances were normalized with Transcripts Per Million (TPM) mapped reads. Differentially expressed genes were identified by DESeq2 ((v1.18.1)) with fold change >2 and Benjamini–Hochberg false discovery rate adjusted P < 0.05. To perform gene set enrichment assay (GSEA), differential expression analysis results of RNA-seq were used to build the pre-ranked gene lists based on signed –log2(fold change). The pre-ranked gene lists were then run using GSEA Java GUI (v2.3.3) with 100 times permutation (http://software.broadinstitute.org/gsea/index.jsp). RNA-seq heatmaps were generated using TPM values of each gene in each replicate (https://biit.cs.ut.ee/clustvis/). To analyze the biological processes of differentially expressed genes in RNA-seq, gene ontology (GO) analysis was performed (https://biit.cs.ut.ee/gprofiler/gost). Full lists of gene ontology analysis are presented in supplementary files xx. To reduce complexity of GO results, GO classification was perform to classify all GO bins into 16 categories (https://www.animalgenome.org/bioinfo/tools/countgo/). To assess the correlation between two sets of RNA-seq samples in a specific gene set, Euclidean distance analysis was used (https://software.broadinstitute.org/morpheus/).

#### Yeast two-hybrid (Y2H) assays

Y2H budding yeast strain (Takara) was cultured with YEPD+adenine overnight at 30°C. Yeast cells were collected OD 0.5 by centrifugation at 3000 rpm for 3 min. Cells were resuspended and washed 2 times with 0.1 M LiAc (in 1x TE buffer). The bait PGBKT7 (0.5 μg) expressing rixosome, Polycomb, and HP1proteins and prey PGADT7 (0.5 μg) vectors were mixed with 10 μg carrier DNA, and further mixed with yeast cells harvested from 10 ml cultures and resuspended in 50 μl 0.1 M LiAc (in 1x TE buffer). DNA-yeast mixture was incubated with 130 μl 40% PEG 4000 for 30 min at 30°C. For transformation, 21 μl DMSO was added and mixed well with the yeast-DNA mixture, followed by heat shock at 42 °C for 20 min. After incubation on ice for 3 min, the cells were pelleted by centrifugation for 3 min at 4°C. The supernatant was then discarded and sterile water was added to resuspend the cells, which were plated on double selective medium SC plates (Trp-, Leu-) for 3 days at 30°C. Colonies were further transferred to quadruple selective medium SC plates (Trp-, Leu-, His-, Ade-) for 3-4 days at 30°C. For spotting assays, cells were incubated overnight in 4 ml double selective SC medium (Trp-, Leu-). The cells were then diluted to OD600 = 1, 1 ml of which was pelleted, washed once with sterilized water, resuspended in 250 μl sterilized water, and transferred to 96 well plates. Three μl of cell suspension from each well was plated on double-selective medium SC plates (Trp-, Leu-) and quadruple-selective medium SC plates (Trp-, Leu-, His-, Ade-) for 4 days.

#### Statistical analysis

Statistical analyses were conducted using GraphPad Prism 8 software. Statistical significance was evaluated using unpaired two-tail student’s *t*-test. All the qRT-PCR and ChIP-qPCR data is represented as mean +/-deviation from three biological replicates.

Hypergeometric distribution was conducted with the online software: https://systems.crump.ucla.edu/hypergeometric/index.php.

#### Software and Algorithms

Software and algorithms used in this study are listed in Table S6.

## Acknowledgements

We thank the Nikon Imaging Center in the Department of Cell Biology at HMS for access to microscopes and advice, members of the Moazed lab for helpful discussions, and Swapnil Parhad, Juntao Yu, Andy Yuan, and Chen Zhou for comments on the manuscript. D.M. is an Investigator of the Howard Hughes Medical Institute.

---

**Fig. S1.**
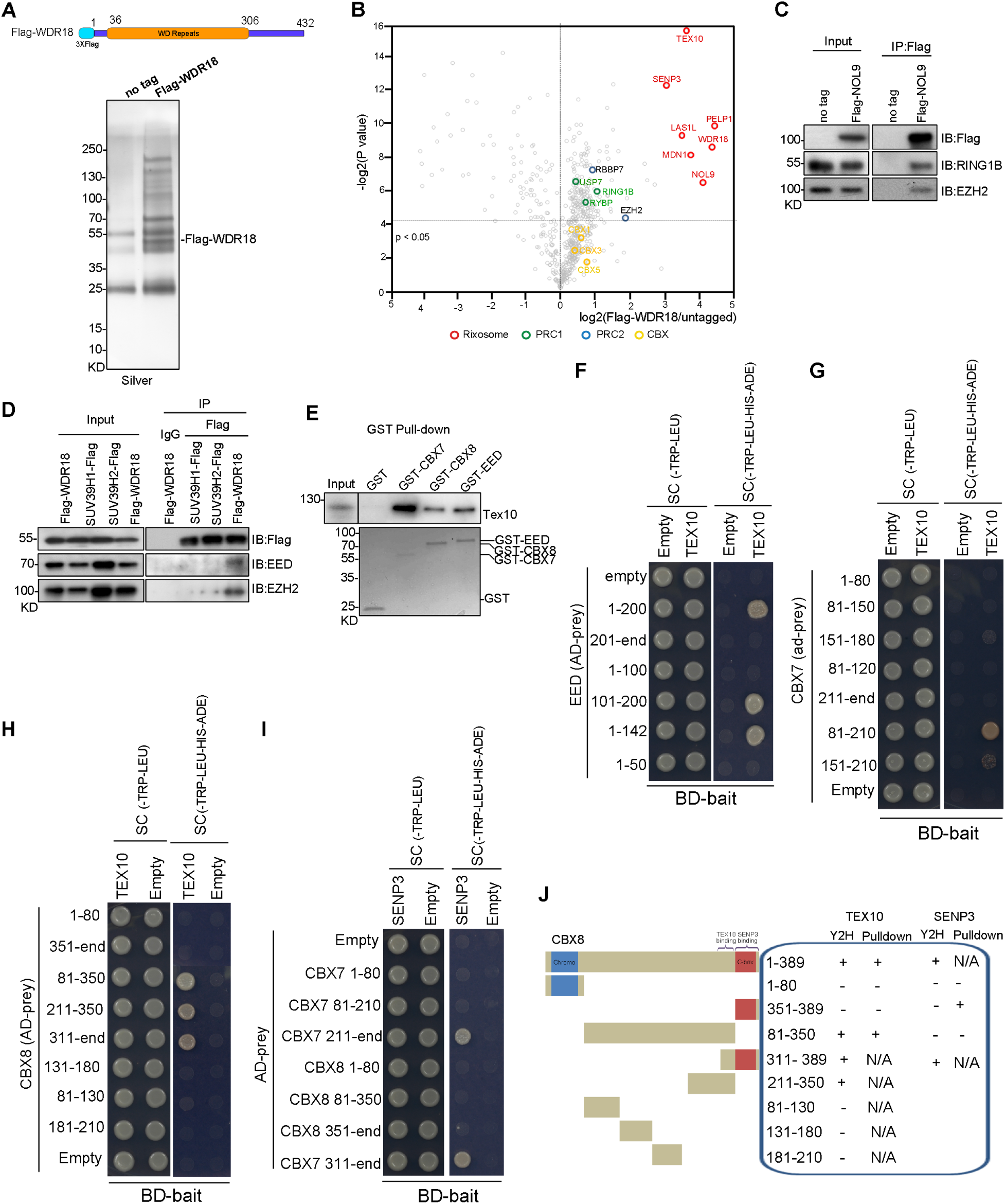
Physical association between the rixosome and Polycomb complexes. (A) Top, schematic diagram for genomic tagging of *NOL9* with 3xFlag, bottom, silver-stained gel of Flag immunoprecipitations from the indicated HEK293FT cell lines. (B) Volcano plot of TMT mass spectrometry results showing log2-fold changes in proteins enrichment in Flag immunoprecipitations from Flag-WDR18 versus untagged cells from two independent experiments. *p* value calculated by t test. Rixosome, PRC1, PCR2, and CBX proteins are highlighted in red, green, blue, and yellow, respectively. (C) Immunoprecipitations (IP) showing the association of RING1B and EZH2 (immunoblot, IB) with Flag-NOL9 in HEK293FT cells. (D) Immunoprecipitations testing associations between the rixosome (Flag-WDR18) and PRC2 subunits (EED, EZH2) or SUV39H1 and SUV39H2. (E) Pull down assays using bacterially expressed and purified TEX10 proteins and bead immobilized GST or the indicated GST-fusion proteins. TEXT10 was detected by immunoblotting using an anti-TEX10 antibody. GST-tagged proteins were stained with Coomassie blue. (F-I) Yeast two-hybrid assays testing interactions between the indicated prey and bait proteins. Yeast cells were transformed with the indicated plasmids and plated onto double selective medium (Trp–, Leu–) or quadruple selective medium (Trp–, Leu–, His–, Ade–). AD, pGADT7; BK, pGBKT7. (J) Schematic diagram summarizing pull-down and Y2H interactions between CBX protein fragments and rixosome subunits (TEX10 or SENP3).

**Fig. S2.**
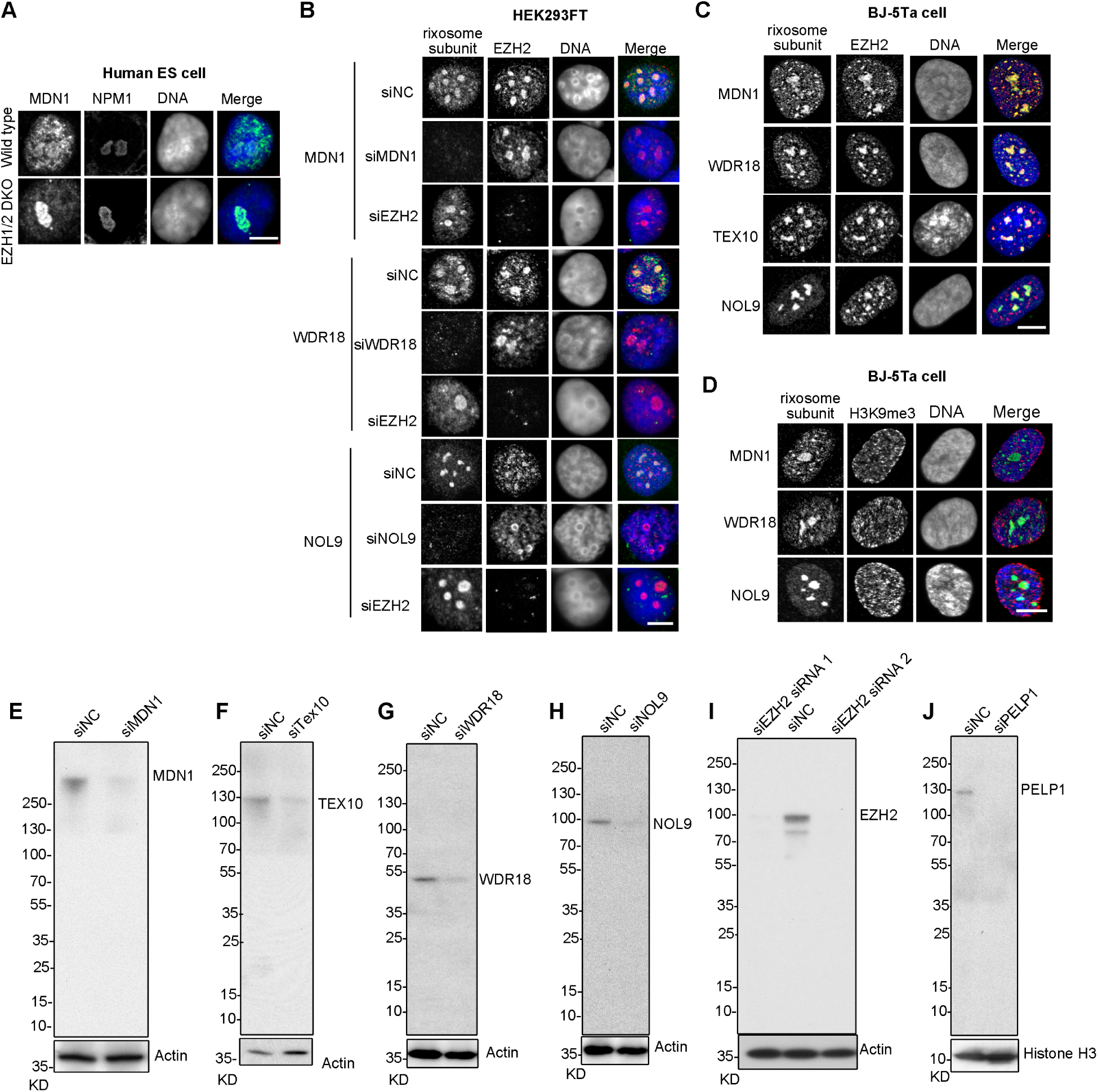
Colocalization of the rixosome with EZH2. **(A)** Immunofluorescence showing the localization of the MDN1 subunit of the rixosome (green) and the nucleolar NPM1 protein (red) in wild-type or *EZH1/2* double knockout (DKO) human ES cells. DNA was stained with DAPI (blue). Scale bar, 5 μm. **(B)** Immunofluorescence of rixosome subunits (MDN1, WDR18, NOL9)(green) and EZH2 (red) in HEK293FT cells treated with the indicated siRNAs. DNA was stained with DAPI (blue). Scale bar, 5 μm. **(C)** Double immunofluorescence of MDN1, WDR18, TEX10, or NOL9 (green), with EZH2 (red) in BJ-5ta cells. DNA was stained with DAPI (blue). Scale bar, 5 μm. **(D)** Double immunofluorescence of MDN1, WDR18, and NOL9 (green), with H3K9me3 (red) in BJ-5ta cells. DNA was stained with DAPI (blue). Scale bar, 5 μm. **(E-J)** Immunoblotting showing protein levels in the siRNA-treated cells detected with the indicated antibodies in HEK293FT cells. Actin and Histone H3 served as a loading control (bottom).

**Fig. S3.**
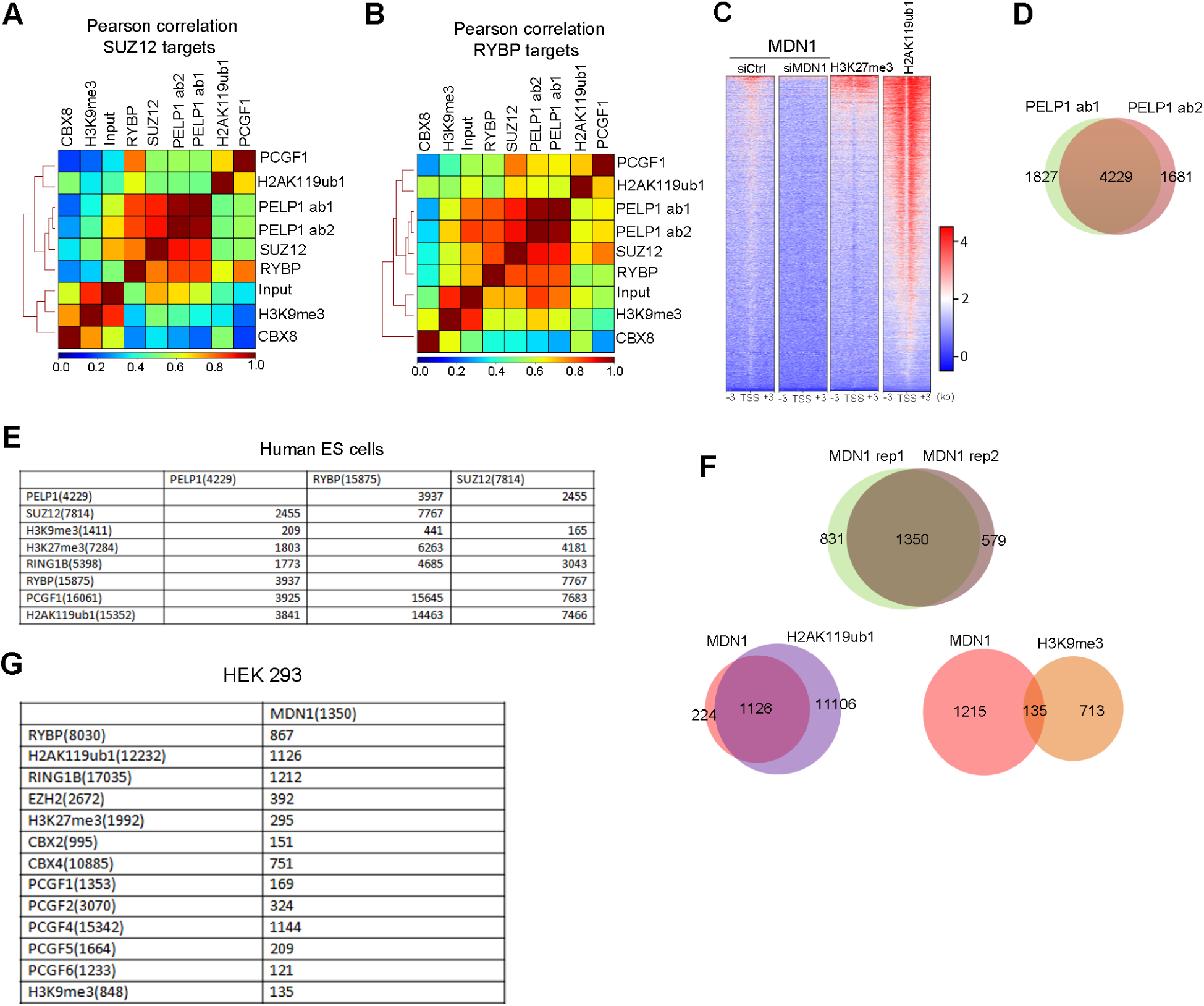
Genomic co-localization of rixosome and Polycomb target loci. **(A, B)** Pearson correlation between SUZ12 (**A**) and RYBP (**B**) enriched regions (based on peak calling of ChIP-seq reads, *n*=15,770 for SUZ12 and 29,433 for RYBP) and regions enriched for the indicated proteins or modifications. (**C**) Heatmaps of MDN1 ChIP-seq localization of MDN1 (siCtrl versus siMDN1), H3K27me3, and H2Ak119ub1 in HEK 293FT cells at transcription start sites (+/-3kb) of annotated RefSeq coding genes (hg19). **(D)** Venn diagram showing overlap between PELP1 ab1 (antibody 2) and PELP1 ab2 (antibody 2) ChIP-seq reads in human ES cells. **(E)** Table showing target gene overlap between the indicated proteins based on ChIP-seq data in human ES cells. **(F)** Venn diagrams showing overlap between indicated ChIP-seq reads in HEK293 cells. Rep, replicate. **(G)** Table summarizing target gene overlaps between MDN1 and the indicated proteins or histone modifications.

**Fig. S4.**
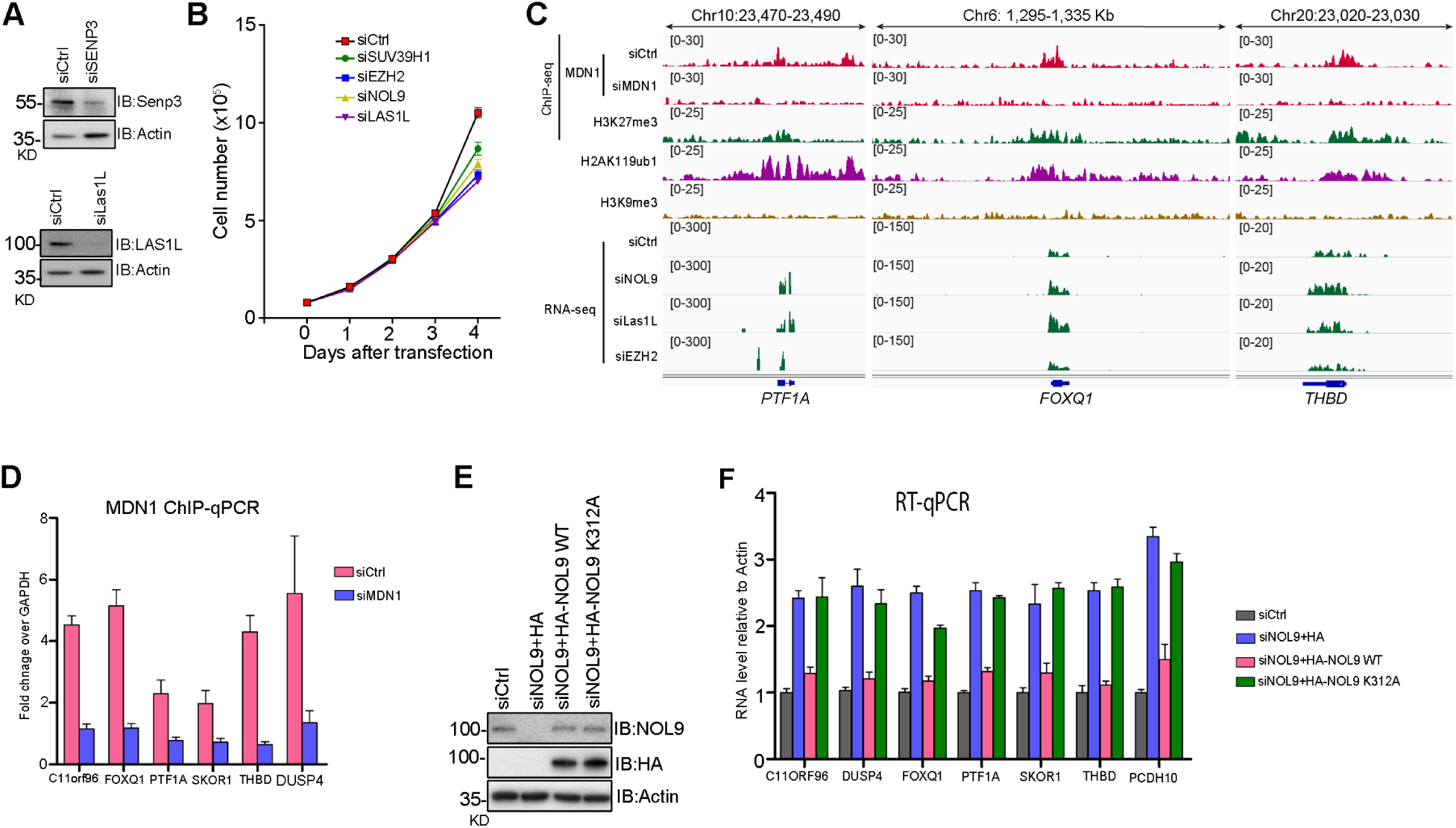
The rixosome is required for repression of target genes. **(A)** Immunoblotting showing the levels of SENP3 subunit of the rixosome control siRNA (siCtrl)- and siSENP3-treated cells. Actin served as a loading control. **(B)** Cell growth curves show cell number changes at indicated time points after knockdowns with siCtrl, siSUV39H1, siEZH2, siNOL9, or si LAS1L siRNA in HEK293 cells. Error bars represent standard deviation for three biological replicates. **(C)** Genomic snapshots of Polycomb target genes *PTF1A,FOXQ1*, and *THBD* showing RNA-seq for the indicated siRNA treatments (siCtrl, siNOL9, siLAS1L, and siEZH2) in HEK293FT cells (bottom 4 tracks). ChIP-seq enrichment of MDN1 (siCtrl and siMDN1), H3K27me3, H2AK119ub1, and H3K9me3 (top 4 tracks). **(D)** ChIP-qPCR showing the localization of MDN1 at the indicated Polycomb target genes in HEK293 cells. siRNA knockdowns (siCtrl and siMDN1) were used to confirm the specificity of the anti-MDN1 antibody. Error bars represent standard deviations for three biological replicates. **(E)** NOL9 knockdown and rescue in HEK293 cells. Immunoblot showing protein levels in control siRNA (siCtrl) and siNOL, and siNol9+NOL9 wild type or NOL9-K312A expressing plasmids. Actin served as a loading control. **(F)** RT-qPCR analysis of RNA levels for the *C11ORF96, DUSP4, FOXQ1, PTF1A, SKOR1, THBD*, and *PCDH10* genes in the indicated siRNA-treated and NOL9-rescued HEK293 cells. RNA expression levels were normalized to housekeeping gene *ACTB*. Error bars represent standard deviations for three biological replicates.

**Fig. S5.**
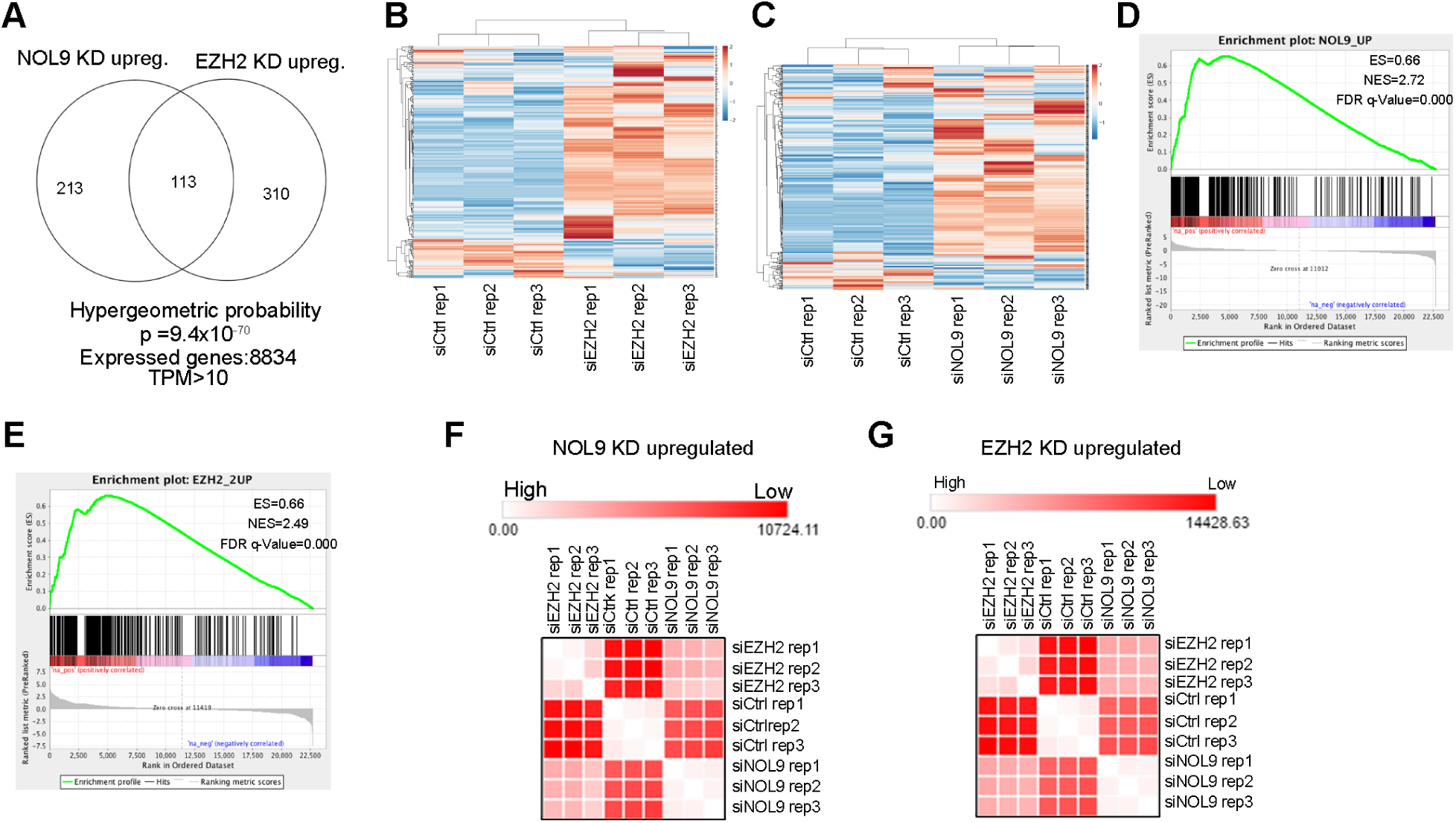
Rixosome represses Polycomb target genes. **(A)** Venn diagram of the overlap between gene sets upregulated in NOL9 KD and EZH2 KD cells. Overlap significance evaluated by calculating hypergeometric probability. Genes with Transcript per Million (TPM)> 10 were considered as expressed. **(B)** Heatmap of RNA-seq expression z-scores in siCtrl and siEZH2 knockdown cells computed for genes that are upregulated in siNOL9 RNA-seq data (*n*=326). **(C)** Heatmap of RNA-seq expression z-scores siCtrl and siNOL9 knockdown cells computed for genes that are upregulated in siEZH2 RNA-seq reads (*n*=423). **(D)** Gene set enrichment analysis of siEZH2 KD RNA-seq results. Gene set is the NOL9 KD upregulated genes (*n*=326). **(E)** Gene set enrichment analysis of siNOL9 KD RNA-seq results. Gene set is the EZH2 KD upregulated genes (*n*=423). **(F)** Euclidean distance analysis showing correlation of log2-fold changes of siNOL9 KD and siEZH2 KD in NOL9 KD upregulated gene set. **(G)** Euclidean distance analysis showing correlation of log2-fold changes of siNOL9 KD and siEZH2 KD in EZH2 KD upregulated gene set.

**Fig. S6.**
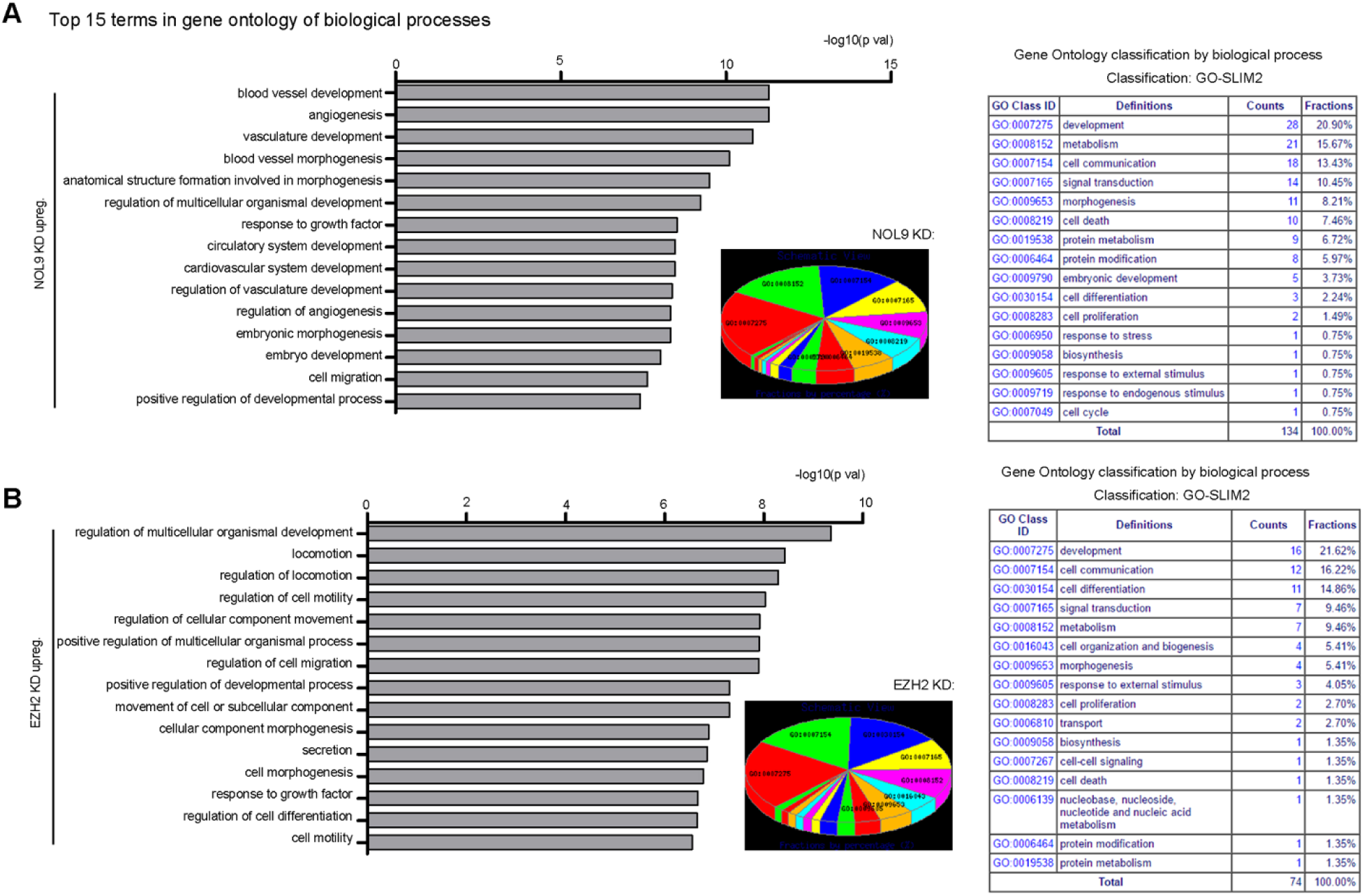
The rixosome and Polycomb pathways represses similar sets of development- and cell differentiation-associated genes. **(A)** Left, gene ontology (GO) analysis showing the biological processes associated with genes upregulated in NOL9 knockdown (KD) HEK293FT cells. A bar plot shows the 15 GO terms (with smallest *p*-values) in GO analysis. A full list of GO terms is classified into 16 classes. Middle, pie chart shows percent of each type of GO classification. Right, table showing GO class identification (ID), definition, counts, and percent of each GO class. **(B)** Left, gene ontology (GO) analysis showing the biological processes associated with genes upregulated in EZH2 KD HEK293 cells. A bar plot shows the 15 GO terms (with smallest *p*-values) in GO analysis. A full list of GO terms is classified into 16 classes. Middle, pie chart shows percent of each type of GO classification. Right, table showing GO class ID, definition, counts, and percent of each GO class.

**Fig. S7.**
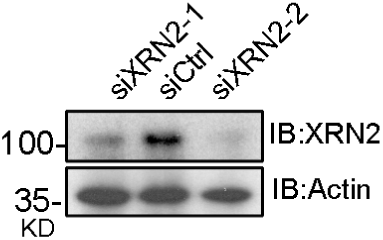
Validation of siXRN2 Knockdown. Immunoblot showing XRN2 protein levels in siCtrl and siXRN2 (−1 and −2 indicate different siRNAs) knockdown cells. Actin served as a loading control.

**Fig. S8.**
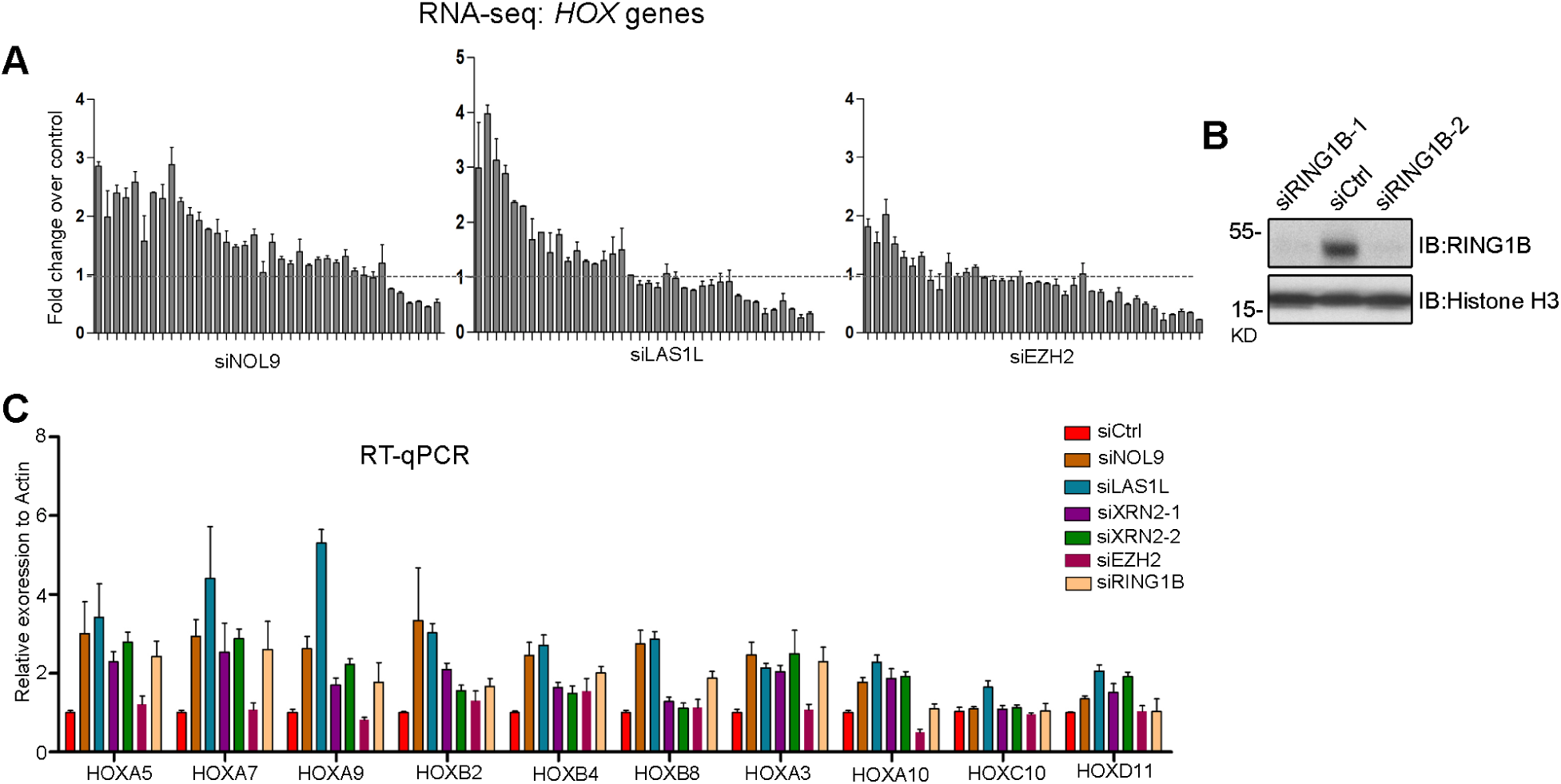
The rixosome targets and represses *HOX* genes. **(A)** Bar plots showing fold changes in the expression (RNA-seq) of 37 *HOX* genes (all except for *HOXB1*) in siNOL9, siLAS1L, or siEZH2 relative to siCtrl knockdowns in HEK293FT cells. **(B)** Immunoblot showing RING1B protein levels in siCtrl versus siRING1B (−1 and −2 indicate different siRNAs) knockdown cells. Actin served as a loading control. **(C)** RT-qPCR showing the effect of the indicated siRNA knockdowns on *HOX* gene expression in HEK293 cells. RNA expression levels were normalized to *ACT1B* mRNA levels. Error bars represent standard deviations for three biological replicates.

**Fig. S9.**
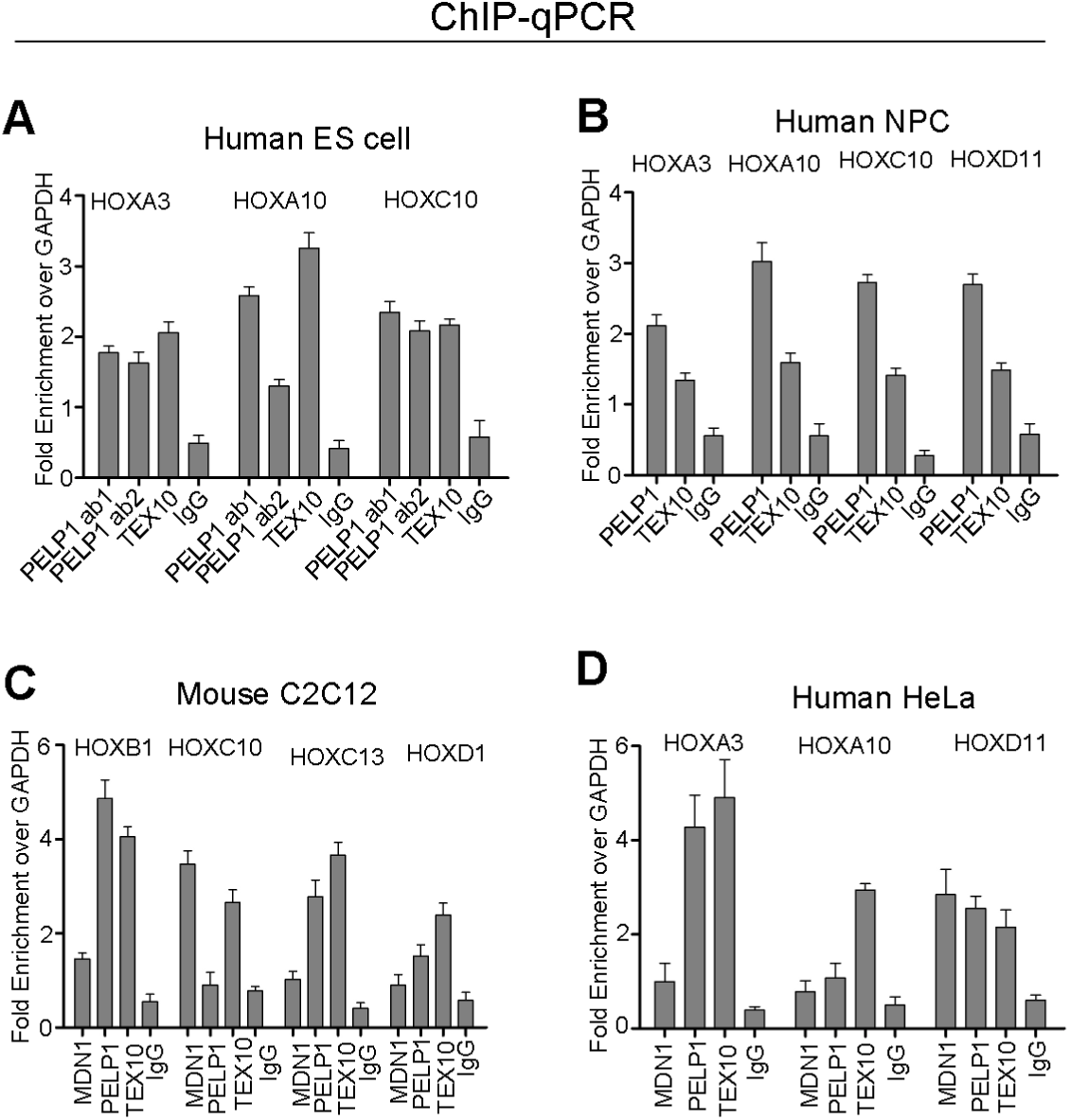
The rixosome associates with Polycomb target genes in various human and mouse cells. **(A-D)** ChIP-qPCR analysis of PELP1 and TEX10 at the indicated *HOX* genes in human ES cells (**A**), human neural progenitor cells, NPCs (**B**), mouse myoblast cells C2C12 (**C**), and human HeLa cells (**D**). Rabbit IgG served as a control. Error bars represent standard deviations for three biological replicates.

**Fig. S10.**
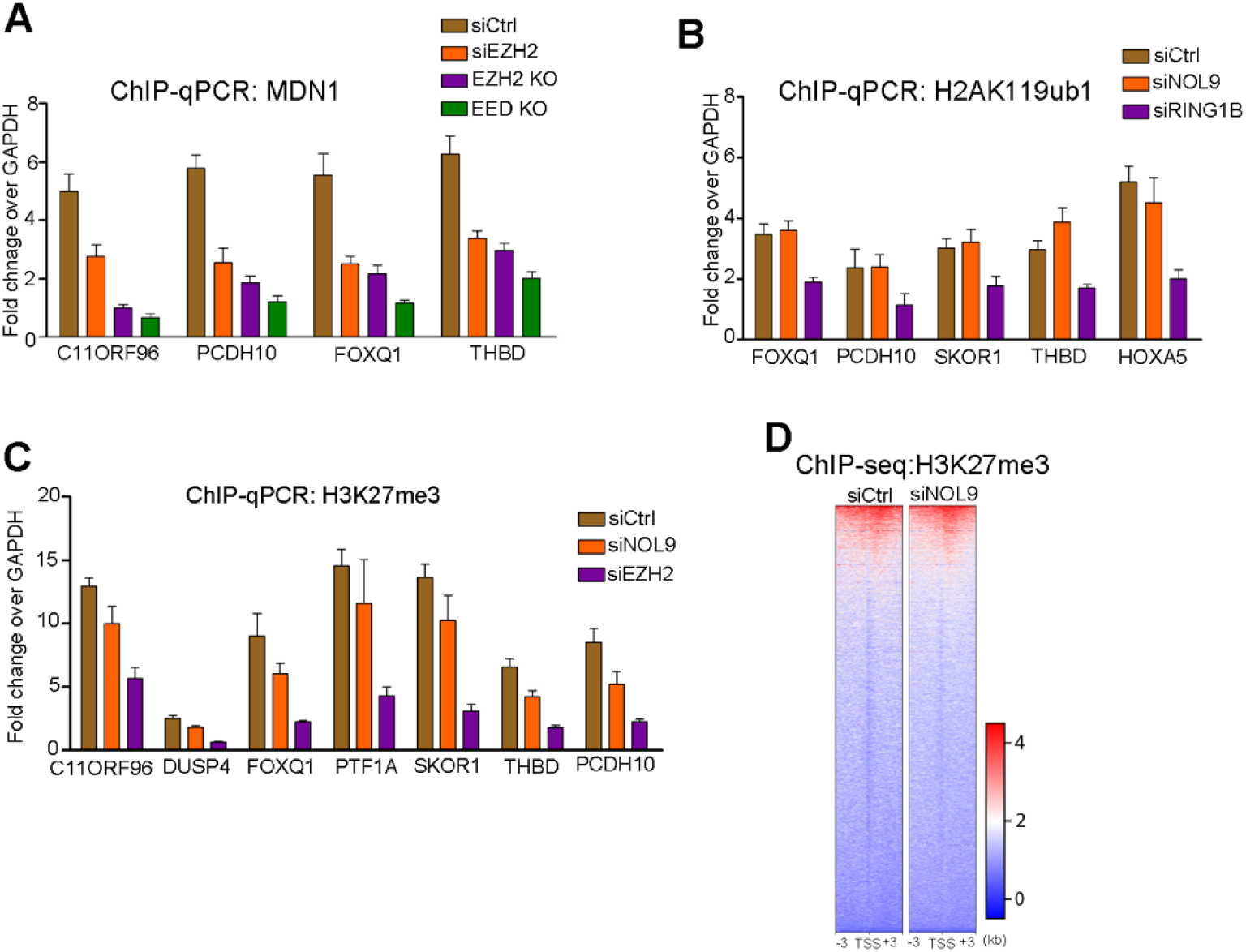
The rixosome is recruited by Polycomb but is not required for H2AK119ub1 or H3K27me3. **(A)** ChIP-qPCR showing the enrichment of MDN1 at the indicated genes in siCtrl, siEZH2, EZH2 knockout, or EED knockout clones of HEK293FT cells. **(B)** ChIP-qPCR showing H2AK119ub1 enrichment at the indicated target genes in siCtrl, siNOL9, and siRING1B treated HEK293FT cells. **(C)** ChIP-qPCR showing the enrichment of H3K27me3 at indicated target genes in siCtrl, siNOL9, and siEZH2 treated HEK293FT cells. Error bars represent standard deviations for three biological replicates (panels **A-C**). **(D)** Heatmaps of H3K27me3 ChIP-seq reads in siCrtrl and siNOL9 treated cells at TSSs (+/-3kb) of annotated RefSeq coding genes (hg19).

## References

Breiling, A., Turner, B. M., Bianchi, M. E. & Orlando, V. 2001. General transcription factors bind promoters repressed by Polycomb group proteins. Nature, 412, 651–5.

Buhler, M., Verdel, A. & Moazed, D. 2006. Tethering RITS to a nascent transcript initiates RNAi- and heterochromatin-dependent gene silencing. Cell, 125, 873–86.

Chen, G., Gulbranson, D. R., Hou, Z., Bolin, J. M., Ruotti, V., Probasco, M. D., Smuga-Otto, K., Howden, S. E., Diol, N. R., Propson, N. E., Wagner, R., Lee, G. O., Antosiewicz-Bourget, J., Teng, J. M. & Thomson, J. A. 2011. Chemically defined conditions for human iPSC derivation and culture. Nat Methods, 8, 424–9.

Dellino, G. I., Schwartz, Y. B., Farkas, G., Mccabe, D., Elgin, S. C. & Pirrotta, V. 2004. Polycomb silencing blocks transcription initiation. Mol Cell, 13, 887–93.

Duboule, D. 2007. The rise and fall of Hox gene clusters. Development, 134, 2549–60.

Francis, N. J., Kingston, R. E. & Woodcock, C. L. 2004. Chromatin compaction by a polycomb group protein complex. Science, 306, 1574–7.

Fromm, L., Falk, S., Flemming, D., Schuller, J. M., Thoms, M., Conti, E. & Hurt, E. 2017. Reconstitution of the complete pathway of ITS2 processing at the pre-ribosome. Nat Commun, 8, 1787.

Grau, D. J., Chapman, B. A., Garlick, J. D., Borowsky, M., Francis, N. J. & Kingston, R. E. 2011. Compaction of chromatin by diverse Polycomb group proteins requires localized regions of high charge. Genes & development, 25, 2210–21.

Holoch, D. & Moazed, D. 2015. RNA-mediated epigenetic regulation of gene expression. Nat Rev Genet, 16, 71–84.

Kowalik, K. M., Shimada, Y., Flury, V., Stadler, M. B., Batki, J. & Buhler, M. 2015. The Paf1 complex represses small-RNA-mediated epigenetic gene silencing. Nature, 520, 248–52.

Lau, M. S., Schwartz, M. G., Kundu, S., Savol, A. J., Wang, P. I., Marr, S. K., Grau, D. J., Schorderet, P., Sadreyev, R. I., Tabin, C. J. & Kingston, R. E. 2017. Mutation of a nucleosome compaction region disrupts Polycomb-mediated axial patterning. Science, 355, 1081–1084.

Margueron, R., Justin, N., Ohno, K., Sharpe, M. L., Son, J., Drury, W. J., 3rd, Voigt, P., Martin, S. R., Taylor, W. R., De Marco, V., Pirrotta, V., Reinberg, D. & Gamblin, S. J. 2009. Role of the polycomb protein EED in the propagation of repressive histone marks. Nature, 461, 762–7.

Margueron, R., Li, G., Sarma, K., Blais, A., Zavadil, J., Woodcock, C. L., Dynlacht, B. D. & Reinberg, D. 2008. Ezh1 and Ezh2 maintain repressive chromatin through different mechanisms. Mol Cell, 32, 503–18.

Margueron, R. & Reinberg, D. 2011. The Polycomb complex PRC2 and its mark in life. Nature, 469, 343–9.

Montavon, T. & Duboule, D. 2013. Chromatin organization and global regulation of Hox gene clusters. Philos Trans R Soc Lond B Biol Sci, 368, 20120367.

Motamedi, M. R., Verdel, A., Colmenares, S. U., Gerber, S. A., Gygi, S. P. & Moazed, D. 2004. Two RNAi complexes, RITS and RDRC, physically interact and localize to noncoding centromeric RNAs. Cell, 119, 789–802.

Navarrete-Perea, J., Yu, Q., Gygi, S. P. & Paulo, J. A. 2018. Streamlined Tandem Mass Tag (SL-TMT) Protocol: An Efficient Strategy for Quantitative (Phospho)proteome Profiling Using Tandem Mass Tag-Synchronous Precursor Selection-MS3. J Proteome Res, 17, 2226–2236.

Nowak, D. E., Tian, B. & Brasier, A. R. 2005. Two-step cross-linking method for identification of NF-kappaB gene network by chromatin immunoprecipitation. Biotechniques, 39, 715–25.

Plys, A. J., Davis, C. P., Kim, J., Rizki, G., Keenen, M. M., Marr, S. K. & Kingston, R. E. 2019. Phase separation of Polycomb-repressive complex 1 is governed by a charged disordered region of CBX2. Genes Dev, 33, 799–813.

Schuettengruber, B., Bourbon, H. M., Di Croce, L. & Cavalli, G. 2017. Genome Regulation by Polycomb and Trithorax: 70 Years and Counting. Cell, 171, 34–57.

Shao, Z., Raible, F., Mollaaghababa, R., Guyon, J. R., Wu, C. T., Bender, W. & Kingston, R. E. 1999. Stabilization of chromatin structure by PRC1, a Polycomb complex. Cell, 98, 37–46.

Shipkovenska, G., Durango, A., Kalocsay, M., Gygi, J. P. & Moazed, D. 2019. An RNA Degradation Complex Required for Spreading and Epigenetic Inheritance of Heterochromatin. bioRxiv doi: https://doi.org/10.1101/870766.

Simon, J. A. & Kingston, R. E. 2013. Occupying chromatin: Polycomb mechanisms for getting to genomic targets, stopping transcriptional traffic, and staying put. Mol Cell, 49, 808–24.

Tatavosian, R., Kent, S., Brown, K., Yao, T., Duc, H. N., Huynh, T. N., Zhen, C. Y., Ma, B., Wang, H. & Ren, X. 2019. Nuclear condensates of the Polycomb protein chromobox 2 (CBX2) assemble through phase separation. J Biol Chem, 294, 1451–1463.

Terranova, R., Yokobayashi, S., Stadler, M. B., Otte, A. P., Van Lohuizen, M., Orkin, S. H. & Peters, A. H. 2008. Polycomb group proteins Ezh2 and Rnf2 direct genomic contraction and imprinted repression in early mouse embryos. Dev Cell, 15, 668–79.

Yu, R., Wang, X. & Moazed, D. 2018. Epigenetic inheritance mediated by coupling of RNAi and histone H3K9 methylation. Nature.

